# Matrix Metalloproteinase Response to Acute Resistance Exercise is Altered in Osteoarthritic Adults with Increased Muscle Collagen and PDGFRα+ Mesenchymal Cells

**DOI:** 10.64898/2026.10.11.757523

**Authors:** A. M. Schweitzer, M. J. Abercrombie, M. D. Fliss, A. Thouli, A. P. Wells, L. Wiens, A. L. Kelso, C. J. Mitchell

## Abstract

**Background:** Knee osteoarthritis (OA) is associated with knee extensor weakness, altered myofibre morphology, reduced satellite cell abundance and fibrillar muscle collagen accumulation in the vastus lateralis (VL). Greater fibrillar muscle collagen is associated with reduced knee extensor strength. Yet its underlying mechanisms remain poorly understood. Satellite cells and PDGFRα+ mesenchymal cells express collagen as well as proteins involved in collagen degradation. The present study assessed knee extensor function and myofibre morphology, satellite cells and PDGFRα+ mesenchymal cells in adults with and without moderate-to-severe knee OA. It also investigated collagen remodeling by assessing how enzymes involved in collagen degradation (matrix metalloproteinases [MMPs]) respond to acute resistance exercise in the VL. This study aimed to better understand the cellular contributors to muscle collagen accumulation and knee extensor weakness in knee OA.

**Methods:** Volunteers with and without moderate-to-severe knee OA had a VL muscle biopsy before and 72-hours following resistance exercise. In VL samples, MMP gene expression (qPCR), protein area fraction (immunofluorescence) and enzyme activity (zymography) were assessed. PDGFRα+ mesenchymal cells, satellite cells, satellite cell-MMP9 colocalization and myofibre morphology were assessed by immunofluorescence. Fibrillar muscle collagen content was quantified by Picrosirius red.

**Results:** The OA group had weaker knee extensors (reduced muscle quality and more fibrillar collagen than controls (12.2 ± 4.8%, 8.4 ± 2.9%, *p* = 0.006). Fibrillar collagen was negatively correlated with isokinetic concentric torque (*r* = -0.479, *p* = 0.006). Volunteers with OA had more PDGFRα+ cells (*p* < 0.001) and fewer satellite cells than controls (*p* = 0.006). In both groups, resistance exercise upregulated *MMP2* (*p* < 0.001), *MMP9* (*p* = 0.001) and *MMP14* (*p* < 0.001) gene expression as well as MMP2 (*p* < 0.001) and MMP9 (*p* = 0.001) enzyme activity. MMP2 and MMP9 protein area fraction increased in response to resistance exercise in controls, but this response was blunted in the knee OA group (age × time interaction, *p* = 0.041). Satellite cell-MMP9 colocalization fold change post-exercise was different between groups (*p* = 0.048). Volunteers with knee OA had smaller myofibres (*p* < 0.001) and greater myofibre shape irregularity (*p* = 0.039), but no differences in thigh lean mass, capillary content or fibre-type distribution.

**Conclusions:** Although exercise stimulated transcriptional and MMP activity responses in both groups, the ECM-specific protein response was impaired in knee OA. Dysregulated ECM remodeling facilitated by MMPs and alterations in PDGFRα+ mesenchymal and satellite cell abundance may contribute to muscle collagen accumulation and knee extensor weakness in knee OA.

## Introduction

Knee osteoarthritis (OA) is a chronic condition characterized by widespread changes within the joint and functional impairment (Noehren et al., 2018; Roos & Arden, 2016; Serrão et al., 2014; Sharma, 2021). Individuals with knee OA exhibit muscle weakness, reduced satellite cell abundance, greater type IIa/x myofibre proportion and muscle collagen accumulation (Callahan et al., 2015; Noehren et al., 2018; Roos & Arden, 2016; Serrão et al., 2014). Collagen accumulation is associated with reduced myofibre radial expansion and lateral force transmission, as well as knee extensor weakness (Azizi et al., 2017; Noehren et al., 2018; Spadoni et al., 2024); therefore, alterations in the skeletal muscle extracellular matrix (ECM) may contribute to reduced muscle quality.

Collagen accumulation occurs when collagen synthesis exceeds collagen breakdown. Mesenchymal cell populations are the primary cell type involved in collagen deposition in the ECM of skeletal muscle but also secrete enzymes responsible for collagen degradation such as matrix metalloproteinases (MMPs) (Joe et al., 2010; Kedlian et al., 2024; Rubenstein et al., 2025; Zanotti et al., 2010). Platelet-derived growth factor receptor-α (PDGFRα) is a cell surface receptor found on mesenchymal cells often classified as fibroadipogenic progenitor (FAP) cells. PDGFRα+ mesenchymal cells are a heterogenous cell population, however, and likely also include fibroblasts (Contreras et al., 2019; Murphy et al., 2011). PDGFRα+ mesenchymal cells have been implicated in muscle collagen accumulation in humans (Cardone et al., 2023; Fry et al., 2017). Yet, there is limited work characterizing the molecular muscle phenotype of individuals with knee OA, including assessment of PDGFRα+ mesenchymal cells and the contribution of blunted collagen breakdown to muscle collagen accumulation.

Our previous work has demonstrated evidence of collagen remodeling 72h after acute resistance exercise in healthy young and older adults, reflected by increased MMP gene expression, protein abundance, and enzyme activity, as well as changes to tissue inhibitor of metalloproteinase (TIMP) gene expression (Schweitzer et al., 2025). MMPs may also contribute to satellite cell niche remodeling, activation, and proliferation (Rayagiri et al., 2018) and may influence muscle adaptation. Resistance exercise is an important therapeutic strategy for knee OA, specifically when there is a goal of avoiding total knee arthroplasty (Roos & Arden, 2016; Sharma, 2021). Therefore, understanding how regulators of collagen breakdown respond to acute resistance exercise in muscle from individuals with knee OA may provide insight into the regulation of ECM remodeling, mechanisms underlying muscle collagen accumulation and how to optimize resistance exercise as a therapeutic. Furthermore, characterizing the cellular and molecular muscle phenotype associated with moderate-to-severe knee OA may provide additional insight into muscle dysfunction in this condition.

The present study aimed to describe how markers of collagen degradation (MMP/TIMP gene expression, MMP protein and activity) respond to acute resistance exercise in the vastus lateralis (VL) from volunteers with and without moderate-to-severe knee OA. It also aimed to assess satellite cells and satellite cell-MMP9 colocalization in the VL both at rest and in response to acute resistance exercise. Further, it aimed to characterize myofibre morphology and capillary content in the VL and investigate the relationships between cellular characteristics and functional strength outcomes. It was hypothesized that compared to controls, volunteers with moderate-to-severe knee OA would have weaker knee extensors and poorer muscle quality, muscle collagen accumulation, more PDGFRα+ mesenchymal cells and less satellite cells, and dysregulated MMP and TIMP response to acute resistance exercise.

## Methods

### Participants

This study was approved by the University of British Columbia Research Ethics Board (H24-03721), and written informed consent was provided by every participant. Twenty participants with moderate-to-severe knee OA and 16 sex- and age-matched controls without knee OA and no history of major knee impairment participated in this study. Participants were age-matched within five years. Due to different group sample sizes, sex-matching was not perfectly even (control, *n* = 8 females, *n* = 8 males; OA, *n* = 14 females, *n* = 6 males). Participants were completing less than two hours per week of lower-body resistance training, had a body mass index (BMI) less than 35, were nonsmokers (cigarettes), were not currently diagnosed with cancer and had not undergone cancer treatment in the past year. They were nondiabetic (type 1 or type 2), non-hypertensive and not on anticoagulants or antiplatelet medication nor had a diagnosis of a bleeding disorder.

### Classification of Knee OA Severity using the Western Ontario and McMaster Universities Osteoarthritis Index (WOMAC)

The present study aimed to recruit a population for whom knee OA had a clinically meaningful impact. Radiographic severity does not always correspond with symptom severity, as individuals with severe radiographic OA may report minimal symptoms, whereas others with relatively minor radiographic changes may experience substantial pain and functional limitations. Therefore, the WOMAC was used to characterize the clinical severity of knee OA.

The WOMAC is a patient-reported outcome measure used to assess symptoms and physical function associated with hip and knee OA (Bellamy et al., 1988). For the present study, participants were instructed to complete the questionnaire with reference to their knee joints specifically. The questionnaire consists of 24 items across three domains: pain (5 items), stiffness (2 items) and physical function (17 items). Each item is scored from 0 to 4, where 0 indicates no symptoms/difficulty and 4 indicates extreme symptoms/difficulty. Results from each item are summed to provide a total score out of 96, with scores also reported separately for each domain. In addition to providing verbal confirmation of a physician-confirmed knee OA diagnosis, participants were required to have a WOMAC score ≥ 40. This threshold was selected to identify participants with a clinically meaningful, moderate-to-severe symptom phenotype, based on evidence from populations undergoing total knee arthroplasty, who have reported mean WOMAC scores of 45.1 ± 15.3 (Kuklinski et al., 2022). There are currently no established WOMAC thresholds for categorizing knee OA severity, however, making the use of a WOMAC score ≥ 40 to characterize a moderate-to-severe phenotype a possible limitation of the present study.

### Study Design

*Visit 1* included completion of the WOMAC (control group only, the OA group completed the WOMAC over the phone during a pre-screening call), anthropometric measurements (height and weight), dual-energy *x*-ray absorptiometry (DXA) and maximal exercise testing. Height and weight were collected by standard methods; volunteers wore minimal indoor clothing and were not wearing shoes. *Visit 2* included a baseline muscle biopsy (0h) and an acute bout of lower body resistance exercise. *Visit 3* took place 72 hours (72h) after visit one and consisted of a muscle biopsy.

Participants were fasted for a minimum of six hours prior to each muscle biopsy and were instructed to refrain from high-intensity exercise outside of the lab for a minimum of 48h before visit two and throughout the 72h between visits two and three. Participants in the OA group who were receiving knee injections were required to wait a minimum of 60 days before scheduling visits two and three to minimize potential local effects of the injection on muscle metabolism. All participants were instructed to avoid taking non-steroidal anti-inflammatory drugs between visits two and three. All muscle biopsies were sampled from the VL under local anaesthesia (1% xylocaine with epinephrine) using a Bergström needle that had been modified for suction (Tarnopolsky et al., 2011). Excess blood, fat and connective tissue were removed from samples, and they were either immediately frozen in liquid nitrogen or embedded in optimal cutting temperature (OCT) compound before freezing in 2-methylbutane cooled by liquid nitrogen. All samples were stored at -80°C for subsequent analysis. In the OA group, muscle biopsies were obtained from the leg affected by OA. For participants with bilateral knee OA, biopsies were obtained from the more symptomatic leg. The control group was randomized to match the distribution of right and left legs sampled in the OA group. All participants were right-leg dominant, and the proportions of biopsies obtained from the dominant and non-dominant legs were similar between groups (control, *n* = 8 right, *n* = 8 left; OA, *n* = 9 right, *n* = 11 left).

### DXA

DXA was used to assess body fat percentage, fat-free, bone-free, lean body mass (LBM) and thigh lean mass (TLM). TLM was calculated from a segmented thigh region of interest (ROI) on the DXA scan. The superior boundary of the ROI was defined by a line extending from the lateral aspect of the anterior superior iliac spine, through the femoral head to the inferior ramus of the pubis. The inferior boundary was defined by a line extending through the centre of the knee joint, parallel to the tibial plateau. The superior and inferior boundaries were connected along the lateral and medial aspects of the limb to encompass the entire thigh.

### Maximal Exercise Testing

During Visit 1, exercise tests were completed in the following order: leg press 10-repetition maximum (10RM), maximal grip strength, knee extensor performance on a Biodex dynamometer and peak oxygen consumption (V̇O_2peak_). Leg press (Atlantis Strength, PE-225, Laval, QC) 10RM was determined within five sets. Weight increased incrementally until the participant was unable to complete 10 repetitions. 1RM was extrapolated from the highest weight achieved and the number of repetitions completed at that weight (Brzycki, 1993). Maximal grip strength was tested using a handgrip dynamometer (Handeful, Digital Hand Dynamometer) following methods described previously (Hoffmann et al., 2019). The sum of the right and left hands was recorded as the total grip strength.

Knee extensor strength and rate of torque development (RTD) were assessed with a computerized dynamometer (Biodex System 4, Biodex Medical, Shirley, NY). Participants were seated with their trunk, hip and working leg secured with straps. Hips were positioned at approximately 90° of flexion, and the backrest was adjusted so that the knee joint extended approximately two finger widths beyond the edge of the seat. Each task was performed bilaterally, starting with the left leg. Knee extensor strength was estimated across three maximal isometric contractions as well as three maximal isokinetic concentric contractions. Isometric contractions were performed at 60° of knee flexion and each contraction was held for three seconds. Isokinetic contractions were performed at 60°/s. Verbal encouragement was kept consistent between participants. Knee extensor RTD was estimated from each isometric contraction, with the peak value recorded from the three trials. Participants were instructed to “generate as much force in as little amount of time as possible” and to be “as powerful as possible” during the isometric contraction trials. The analog torque signal was captured using an external analog-digital convertor (ADInstruments, Colorado Springs, CO) at a sampling rate of 2000 Hz and analyzed on LabChart software (ADInstruments, Colorado Springs, CO). Raw data were converted to Nm using a conversion factor calculated from the known torque of the Biodex calibration arm. A low-pass 10 Hz filter was applied to the data, after which a secondary channel was created, and the first derivative of the filtered signal was calculated. From the first derivative channel, the onset of force generation was identified and RTD was recorded as the peak value in the subsequent 200ms. The Biodex data presented in the results section correspond to the biopsied leg.

To assess V̇O_2peak_, volunteers completed an incremental ramp test to exhaustion on a cycle ergometer (Monark, LC6, Sweden). All participants completed a five-minute warm-up at 20 W. After the warm-up, resistance gradually increased at a rate of 1 W every 3-6 seconds. The work rate was adjusted based on the participant’s sex and body size. The test was stopped if the participant indicated they had reached volitional fatigue or if they could no longer maintain a cadence of at least 60 RPM. V̇O_2peak_ was accepted as the highest 30-second average V̇O_2_ recorded during the test. V̇O_2peak_ percentiles were extrapolated from normative datasets (Kaminsky et al., 2022).

### Acute Resistance Exercise

Immediately following the baseline muscle biopsy, participants completed a warm-up on the leg press machine (Atlantis Strength, PE-225, Laval, QC) that would be used for their acute exercise bout. The warm-up consisted of 1 set of 8 repetitions at a very low load, followed by 1-2 sets of 4-6 repetitions at a moderate load, building towards their working load of 70% 1RM. Before starting their first exercise set, participants completed one final warm up of 2-4 repetitions at their working load. They then completed eight sets of leg press (Atlantis Strength, PE-225, Laval, QC) to exhaustion at 70% 1RM. The number of repetitions was maintained between 8 and 20, with the load adjusted between sets as necessary.

### Picrosirius Red

Muscle pieces frozen in OCT compound were sectioned at 10 μm thickness on a cryostat (HM 525NX, Thermo Scientific, Waltham, MA) and equilibrated to room temperature before staining. Slides were bathed in xylene (MilliporeSigma, Burlington, MA) for 30 minutes, allowed to dry completely and rehydrated for five minutes each in 100%, 90%, 80% and 70% ethanol. After a quick wash in distilled water, slides were immersed in Picrosirius red solution (100 mL picric acid, 1.2% Ricca Chemical, Arlington, TX, 0.1% direct red, Thermo Scientific, Waltham, MA) for 45-minutes. Afterward, slides were rinsed quickly in distilled water and immersed in 0.5% glacial acetic acid (Fisher Chemical, Pittsburgh, PA) for two minutes. Slides were then dehydrated for one minute each in 70%, 80%, 90% and 100% ethanol. After drying completely, slides were immersed in xylene for one hour, allowed to dry once again and mounted with Cytoseal XYL (Epredia, Kalamazoo, MI). Images were captured at ×10 magnification on an Evos FL Auto 2 Microscope (Thermo Scientific, Waltham, MA) and analysis was completed on ImageJ software (National Institutes of Health, Bethesda, MD). The analysis was completed in a blinded manner. Using the thresholding features of ImageJ the total fibrillar collagen between myofibres, relative to the total muscle area, was measured and reported as a percentage.

### Immunofluorescence Staining Procedures

Muscle pieces frozen in OCT compound were serially sectioned at 5 μm thickness on a cryostat (HM 525NX, Thermo Scientific, Waltham, MA) with no more than 10 μm between serial sections and stored at -80°C until use. Samples were equilibrated to room temperature before staining. Serial sections were stained with the following antibody panels: *1)* Pax7/MMP9/laminin/4’,6-diamidino-2-phenylindole, dihydrochloride (DAPI), *2)* myosin heavy chain (MHC) (slow fibres)/laminin/DAPI, *3)* PDGFRα/MMP2/laminin/DAPI and, *4)* neural cell adhesion molecule (NCAM)/laminin/DAPI (Table 1).

**Table 1:** Immunofluorescence Antibody Panels on Serial Sections. DAPI, 4’,6-diamidino-2-phenylindole; MMP2, matrix metalloproteinase-2; MMP9, matrix metalloproteinase-9; MHC, myosin heavy chain; NCAM, neural cell adhesion molecule; Pax7, paired box-7; PDGFRα, platelet-derived growth factor receptor-alpha.

| Panel | Antibody 1 | Antibody 2 | Antibody 3 | Nuclear Stain |
| --- | --- | --- | --- | --- |
| 1 | Pax7 | MMP9 | Laminin | DAPI |
| 2 | MHC (slow fibres) | Laminin |  | DAPI |
| 3 | PDGFRα | MMP2 | Laminin | DAPI |
| 4 | NCAM | Laminin |  | DAPI |

NCAM frozen cells (DSHB, 5.1H11) were sent to the University of British Columbia Antibody Proteomics and Biologics Core for purification before dilution. Appropriate secondary antibodies were used for immunofluorescent detection. Detailed antibody information is found in Table 2.

**Table 2:** Immunofluorescence Antibody Information. MMP2, matrix metalloproteinase-2; MMP9, matrix metalloproteinase-9; MHC, myosin heavy chain; NCAM, neural cell adhesion molecule; Pax7, paired box-7; PDGFRα, platelet-derived growth factor receptor-alpha.

| Primary Antibody | Species | Source | Catalogue Number | Dilution |
| --- | --- | --- | --- | --- |
| Anti-laminin | Rabbit | Abcam | ab11575 | 1:500 |
| Anti-Pax7 | Mouse | DSHB | Pax7 | 1:1 |
| Anti-MHC (slow fibres) | Mouse | DSHB | A4.951 | 1:1 |
| Anti-MMP9 | Rabbit | Abcam | ab283575 | 1:50 |
| Anti-MMP2 | Rabbit | Abcam | ab92536 | 1:250 |
| Anti-PDGFRα | Mouse | Bio-Techne | NBP2-52528 | 1:250 |
| Anti-NCAM | Mouse | DSHB | 5.1H11 | 1:500 |
| **Secondary Antibody** |  |  |  |  |
| Alexa Fluor 488 goat anti-mouse |  | Abcam | ab150113 | 1:500 |
| Alexa Fluor 555 goat anti-rabbit |  | Abcam | ab150078 | 1:500 |
| Alexa Fluor 647 goat anti-rabbit |  | Abcam | ab150079 | 1:500 |

For co-immunofluorescence staining, sections for panels *1-3)* were fixed with 4% paraformaldehyde (PFA) (Life Technologies, Waltham, MA) and panel *4)* was fixed in 1% PFA for 10 minutes at room temperature. PFA was rinsed off with three, five-minute washes (3 × 5 minutes) of 1× phosphate buffered saline (PBS) placed directly on the slide. Panel *1)* was blocked in 5% normal goat serum (NGS) (Invitrogen, Waltham, MA), 0.2% Triton X-100 (Fisher Bioreagents, Waltham, MA), 2% bovine serum albumin (BSA) (VWR, Radnor, PA) and 0.15% sodium azide (Thermo Scientific, Waltham, MA) in 1× PBS for 90-minutes at room temperature. Panel *2)* was blocked in 5% NGS and 0.3% Triton X-100 in 1× PBS for 90 minutes at room temperature. Panel *3)* was blocked in 5% NGS, 0.3% Triton X-100 and 1% BSA in 1× PBS-T (1× PBS with 0.1% Tween 20, VWR, Radnor, PA) for one hour at room temperature. Panel *4)* was blocked in 5% NGS, 0.01% Triton X-100 and 1% BSA in 1× PBS-T for one hour at room temperature. Primary antibody cocktails (Pax7 + MMP9; MHC (slow fibres) + laminin; PDGFRα + MMP2; NCAM + laminin) were incubated overnight at 4°C in hydration chambers to prevent evaporation. After overnight incubation, panels *1)* and *2)* were washed 3 × 5 minutes in 1× PBS-T and panels *3)* and *4)* were washed 3 × 5 minutes in 1× PBS. Sections were then incubated in appropriate secondary antibodies for two hours (Panels 1-3) or 90 minutes (Panel 4) at room temperature. Once secondary antibodies were added, all steps following took place protected from light. After secondary incubations, 3 × 5-minute 1× PBS/PBS-T washes were repeated. Panels *1)* and *3)* were then re-fixed in 4% PFA for five minutes at room temperature and 3 × 5-minute 1× PBS/PBS-T washes were repeated. Panels *1)* and *3)* then incubated with anti-laminin primary antibody for two hours at room temperature. After incubation, 3 × 5-minute 1× PBS/PBS-T washes were repeated, and the appropriate secondary antibody was added. Secondaries incubated for two hours at room temperature, followed by 3 × 5-minute 1× PBS/PBS-T washes. Nuclei were labelled with DAPI (1 μg/mL) following manufacturer’s instructions (Invitrogen, Waltham, MA) for five minutes at room temperature. Sections were rinsed with 1× PBS-T, then 1× PBS and cover-slipped with Cytoseal XYL (Epredia, Kalamazoo, MI).

### Immunofluorescence Analyses

Table 3 shows the characteristics of the immunofluorescence analyses.

**Table 3:** Characteristics of Immunofluorescence Analyses. Data are presented as mean ± standard deviation. MMP2, matrix metalloproteinase-2; MMP9, matrix metalloproteinase-9; PDGFRα, platelet-derived growth factor receptor-alpha.

|  | Control |  | OA |  |
| --- | --- | --- | --- | --- |
| **Fibre Number for Morphology/Fibre-Typing Analyses** | 199.2 ± 104.1 |  | 214.4 ± 98.6 |  |
| **Area Assessed for MMP2 Protein Content (mm<sup>2</sup>)** |  |  |  |  |
| Baseline (0h) | 1.4 ± 0.9 |  | 1.1 ± 0.6 |  |
| Post-Exercise (72h) | 1.5 ± 1.3 |  | 1.1 ± 0.6 |  |
| **Area Assessed for MMP9 Protein Content (mm<sup>2</sup>)** |  |  |  |  |
| Baseline (0h) | 1.2 ± 1.2 |  | 0.8 ± 0.5 |  |
| Post-Exercise (72h) | 1.0 ± 0.8 |  | 0.6 ± 0.4 |  |
| **Area Assessed for PDGFRα+ Cell Analysis (mm<sup>2</sup>)** |  |  |  |  |
| Baseline (0h) | 1.8 ± 0.8 |  | 1.5 ± 0.9 |  |
| Post-Exercise (72h) | 1.7 ± 1.3 |  | 1.4 ± 0.8 |  |
| **Fibre Number for Satellite Cell Analysis** | Type I | Type II | Type I | Type II |
| Baseline (0h) | 66.8 ± 35.5 | 69.6 ± 26.0 | 82.3 ± 45.8 | 66.6 ± 33.8 |
| Post-Exercise (72h) | 66.4 ± 20.0 | 61.8 ± 43.6 | 64.7 ± 26.1 | 68.8 ± 38.3 |

Images were captured at ×10 and ×20 magnification (Panels *1* and *3*) or ×10 magnification (Panel *2* and *4*) on an Evos FL Auto 2 Microscope (Thermo Scientific, Waltham, MA). Due to the sensitivity of the thresholding analysis planned for Panels *1)* and *3)*, light, exposure and gain settings were kept constant between all slides during ×10 imaging. All image analysis was completed on ImageJ software (National Institutes of Health, Bethesda, MD) and was done in a completely blinded manner; the assessor did not know the group or time point.

***Panel 1) (×20 magnification)*** Satellite cells were defined as Pax7 colocalized to DAPI beneath the myofibre basal lamina (Nederveen et al., 2016). Incarcerated satellite cells were defined as the lamina covering ≥ 90% of the perimeter of the Pax7+/DAPI+ nucleus (Nederveen et al., 2020). An ROI of 100 myofibres was counted on each sample, and satellite cells were quantified within that region. In conjunction with the fibre-typing of panel *2)*, satellite cell content per type I and type II fibre was quantified. Capillary content per fibre was quantified with the laminin staining from the same 100 myofibre ROI that satellite cells were quantified from. Laminin and DAPI channels were overlaid, and small, round, laminin-positive structures lacking a DAPI-positive nucleus were identified as capillaries. Colocalization of MMP9 and satellite cells was assessed and satellite cells were considered MMP9+ if MMP9 antibody colocalized with Pax7 (Figure 1).

**Figure 1:**
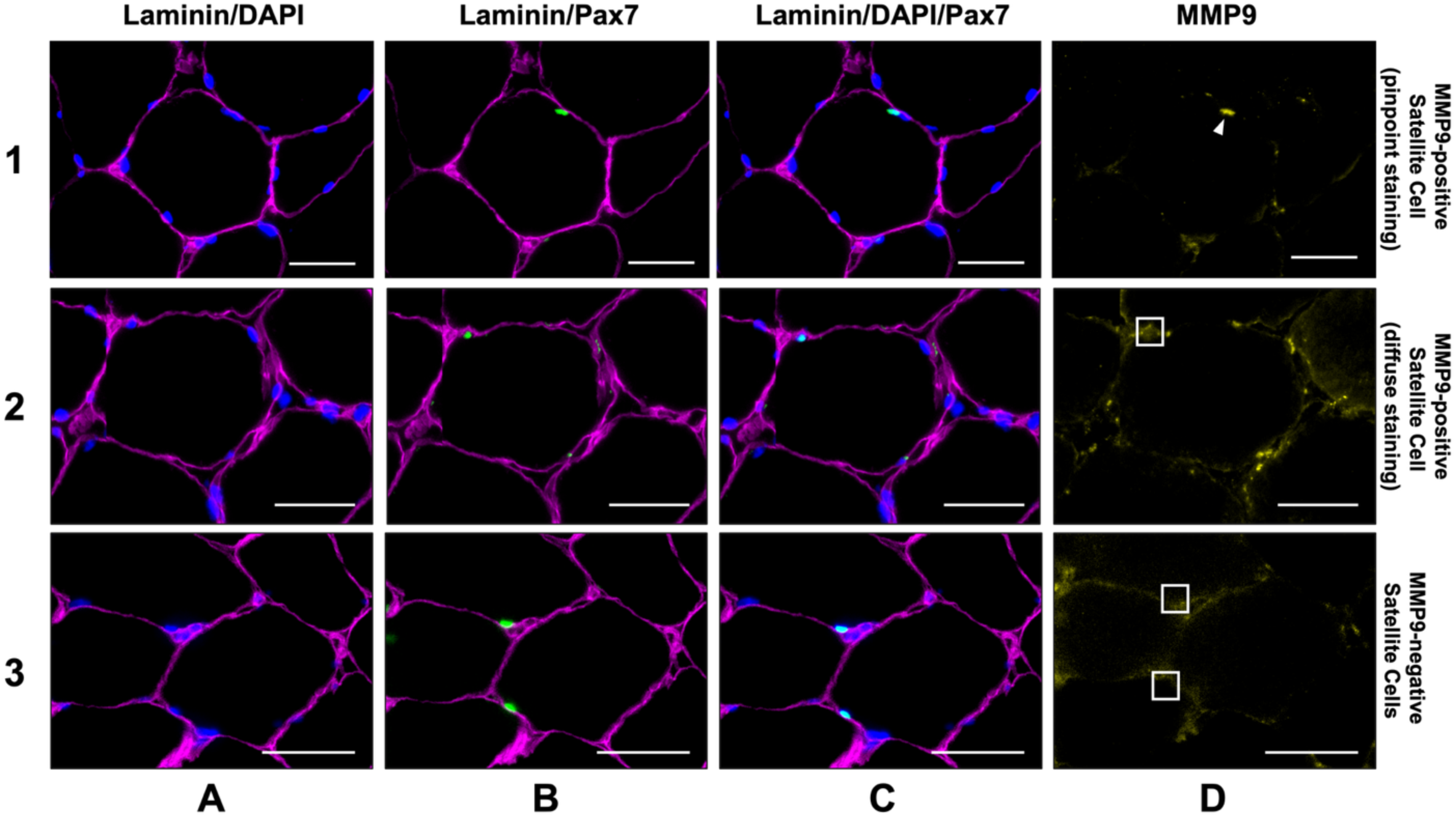
Representative Images of MMP9-positive and MMP9-negative Satellite Cells. *Row 1)* Representative images of MMP9-satellite cell colocalization in a pinpoint fashion. White arrowhead shows MMP9 staining in a pinpoint manner at the location of the satellite cell. *Row 2)* Representative images of MMP9-satellite cell colocalization in diffuse manner. White box shows where MMP9 stains in a diffuse manner around the satellite cell. *Row 3)* Representative images of MMP9-negative satellite cells. White boxes show MMP9-negative area where satellite cells are found. *Column A)* Laminin (magenta) + DAPI (blue). *Column B)* Laminin (magenta) + Pax7 (green). *Column C)* Laminin (magenta) + DAPI (blue) + Pax7 (green), DAPI + Pax7 overlay (cyan). *Column D)* MMP9 (yellow). Scale bar = 50 μm. MMP2, matrix metalloproteinse-2; MMP9 matrix metalloproteinase-9.

***Panel 2) (×10 magnification)*** Fibre size analysis was completed on the baseline muscle biopsy samples by hand using the ROI function on ImageJ (National Institutes of Health, Bethesda, MD). Fibre type distribution was assessed by hand using the mean grey value of the ROIs created during fibre size analysis and the histogram function on ImageJ. Myofibre shape was assessed using the shape factor index (SFI), where values above 1.0 show deviation from a circle (Soendenbroe et al., 2024). Since myofibres are not naturally perfect circles, they have SFIs greater than 1.0, but higher values indicate greater shape deviation (Soendenbroe et al., 2024). SFI was calculated using the ROIs created during fibre size analysis with the following formula:

***Panel 3) (×20 magnification)*** PDGFRα+ mesenchymal cells were defined by PDGFRα staining surrounding the entire nucleus (100% circumferential staining) and localization within the extracellular space (Joe et al., 2010). PDGFRα+ mesenchymal cells were assessed per mm^2^ tissue as has been reported previously (Cardone et al., 2023; Fry et al., 2017). Mesenchymal cells are a heterogeneous cell type and PDGFRα+ cells have numerous subpopulations (Binet et al., 2023; Farup et al., 2021; Rubenstein et al., 2025; Stec et al., 2025) as well as considerable overlap with connective tissue fibroblasts (Contreras et al., 2019; Murphy et al., 2011). As PDGFRα was the cell marker used to assess mesenchymal cells in the present study, the cell population is referred to as PDGFRα+ mesenchymal cells herein.

***Panels 1) and 3) (×10 magnification)*** Using the thresholding features of ImageJ, MMP2 and MMP9 protein area were quantified relative to the total muscle area and were reported as a percentage. Upper and lower thresholds set on ImageJ were kept constant between all MMP2-stained images and all MMP9-stained images.

***Panel 4) (×10 magnification)*** NCAM+ myofibres were counted in a subset of samples (*n* = 10 control; *n* = 17 OA). When NCAM+ staining was present, fibres were counted in the sample, and NCAM+ staining was expressed as a percentage of the total fibres assessed.

### qPCR

RNA was extracted from 15-25 mg of flash-frozen muscle tissue. Tissue was kept cold and homogenized in a well plate homogenizer (Bead Ruptor 96, OMNI International, Kennesaw, GA). After homogenization, 1 mL of 4°C Trizol (Thermo Scientific, Waltham, MA) was added to each sample and incubated on a rocker for five minutes at room temperature. Afterwards, 200 μL of chloroform (Fisher Chemical, Pittsburgh, PA) was added to each sample, and tubes were shaken by hand to mix. Samples were then incubated at room temperature for 3-5 minutes.

Samples were transferred to RNAse-free microcentrifuge tubes, mixed thoroughly with a pipette and centrifuged for 20 minutes at 10000 *g* at 4°C. After centrifugation, the upper aqueous phase was transferred to a new microcentrifuge tube, and the RNA was extracted using the Qiagen RNeasy Kit following manufacturer’s instructions (Hilden, NRW, Germany). cDNA was synthesized from RNA using a high-capacity cDNA reverse transcriptase kit following manufacturer’s instructions (Applied Biosystems, Waltham, MA). For *MMP9* TaqMan gene expression analysis, 80 ng of cDNA was loaded per well in triplicate (Hs00957562) with VCP (Hs00997642) duplexed for a housekeeping gene following manufacturer’s instructions (Applied Biosystems, Waltham, MA). For *MMP2, MMP14, TIMP1* and *TIMP2* SYBR Green (Applied Biosystems, Waltham, MA) gene expression analysis, 10 ng of cDNA was loaded per well in triplicate. *GAPDH* was used as a housekeeping control for all SYBR analyses. See Table 4 for primer sequences; all primers were purchased from Thermo Scientific (Waltham, MA).

**Table 4:**
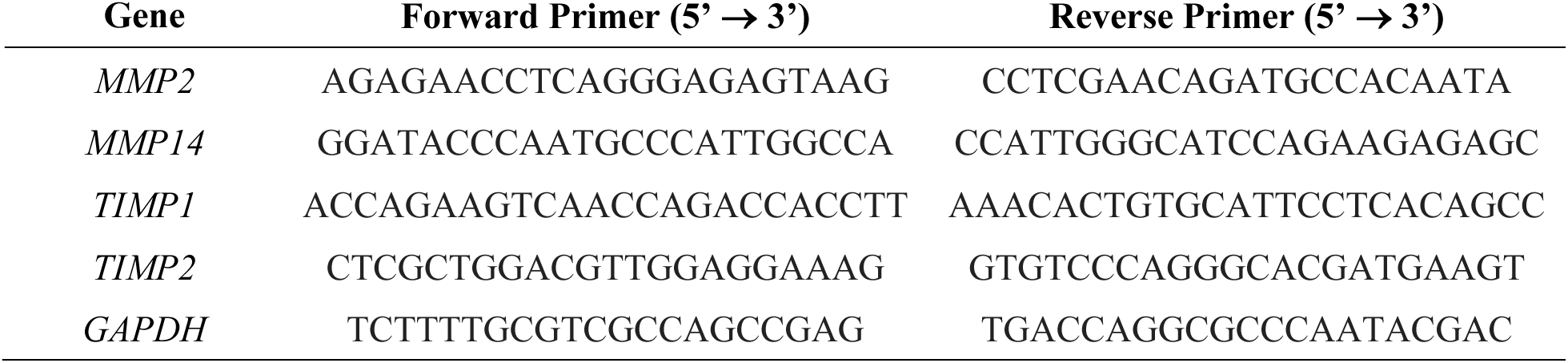
qPCR Primer Sequences. GAPDH, glyceraldehyde 3-phosphate dehydrogenase; MMP2, matrix metalloproteinase-2; MMP14, matrix metalloproteinase-14; TIMP1, tissue inhibitor of metalloproteinases-1; TIMP2, tissue inhibitor of metalloproteinases-2.

### Substrate Zymography

Substrate zymography was used to assess MMP2 and MMP9 enzyme activity potential in frozen tissue. Frozen muscle samples were homogenized in 10 μL/mg RIPA lysis buffer using a well plate homogenizer (Bead Ruptor 96, OMNI International, Kennesaw, GA) and centrifuged at 700 *g* for 15 minutes at 4°C. One complete protease and phosphatase inhibitor mini tablet (Thermo Scientific, Waltham, MA) was added per 10 mL of lysis buffer. The supernatant (sarcoplasmic fraction) was used for subsequent analysis. Total protein was measured via BCA protein assay kit (Thermo Scientific, Waltham, MA) following manufacturer’s instructions. Samples were prepared in 2× SDS zymogram buffer and 4 μg total protein was loaded per well of each 1.0 mm gel (10% acrylamide, 0.5 mg/mL gelatin). Electrophoresis was run at 130 V for 80 minutes. Following separation, gels were incubated in renaturation buffer (2.5% Triton X-100) for 30 minutes at room temperature with gentle agitation. Renaturation buffer was decanted and gels were rinsed in development buffer (50 mM Tris Base, 200 mM sodium chloride, 5 mM calcium chloride, 0.02% Brij-30) for 30 minutes at room temperature with gentle agitation. The development buffer was discarded, and gels were then incubated in fresh development buffer for 15 hours at room temperature.

Through previous optimization experiments, it was determined that zymogram enzyme digestion is sensitive to time since lysate preparation, temperature during development and development time. Therefore, these conditions can be optimized for each zymography experiment. After the second development, gels were stained with 0.5% Coomassie blue in 40% methanol, 10% glacial acetic acid solution for 45 minutes at room temperature with gentle agitation. Following staining, gels were destained in 40% methanol, 10% glacial acetic acid solution for one hour at room temperature with gentle agitation and destaining buffer was changed every 10 minutes. Gels were imaged on a white tray in trans-white light to visualize areas of digestion (indicating areas of greater enzyme activity). Areas of enzyme digestion were evident by a lack of blue staining on the Coomassie-stained gel. Total enzyme activity was quantified using ImageJ software.

During zymography, interactions between MMPs and endogenous inhibitors are disrupted and latent proenzymes may be activated under the conditions of the assay (Snoek-van Beurden & Von den Hoff, 2005). As a result, the activity measured on the zymogram represents the total MMP activity potential within the tissue rather than MMP activity that was necessarily present at the time of tissue collection.

### Statistical Analyses

All statistical analyses were completed using GraphPad Prism version 11.0.2 for Mac OS X (GraphPad Software, Boston, MA). For participant characteristics, normally distributed data were analyzed using Welch’s *t*-test, whereas non-normally distributed data were analyzed by Mann-Whitney test. Knee extensor performance, percent collagen area and capillaries per fibre were analyzed by Welch’s *t*-test. Simple linear regression was used to assess the correlarions between measures of knee extensor performance, percent collagen area, SFI and PDGFRα+ mesenchymal cells. The Pearson correlation coefficient (*r*) and corresponding *p*-value were reported, with statistical significance accepted at *p* < 0.05.

Myofibre morphology and fibre type distribution were analyzed using a 2way ANOVA with Šídák’s multiple comparisons test. ROUT (Q = 1%) outlier test was performed and one outlier was removed from the OA CSA data. qPCR was analyzed using the ΔΔCq method, with the mean of the control-0h time point used as the reference value. Statistical analysis was performed using a mixed-effects model with Šídák’s multiple comparisons test. Outliers were identified using the ROUT method (Q = 1%) and excluded from analysis as follows: *MMP2* (*n* = 1 control; *n* = 1 OA), *MMP9* (*n* = 2 control; *n* = 1 OA), *MMP14* (*n* = 2 control), *TIMP1* (*n* = 2 control; *n* = 1 OA), *TIMP2* (*n* = 2 control). MMP2 and MMP9 percent area were analyzed with a mixed-effects model and Šídák’s multiple comparisons test. Outliers were identified using the ROUT method (Q = 1%) and excluded from analysis as follows: MMP2 (*n* = 1 OA), MMP9 (*n* =

1 control; *n* = 3 OA). Due to between-gel variability, zymography data were analyzed by normalizing the 72h time point to the individuals 0h time point and reported as a FC. Statistical analyses were performed using a mixed-effects model with Šídák’s multiple comparisons test where baseline was set to 1. Outliers were identified using the ROUT method (Q = 1%) and excluded from analysis as follows: MMP2 (*n* = 2 control), MMP9 (*n* = 2 OA). PDGFRα+ mesenchymal cells per mm^2^ were analyzed using a mixed effects model and Šídák’s multiple comparisons test. Outliers were identified using the ROUT method (Q = 1%) and three outliers were removed from the control data and one outlier was removed from the OA data. Satellite cell abundance was analyzed using a mixed-effects model with time, group and fibre type as fixed effects, followed by Tukey’s multiple comparison’s test. Incarcerated satellite cell content was analyzed by 2way ANOVA with Šídák’s multiple comparisons test. Outliers were identified using the ROUT method (Q = 1%) and two outliers were removed from the control data and three outliers were removed from the OA data.

MMP9-satellite cell colocalization percentages were analyzed using a mixed effects model and Šídák’s multiple comparisons test. MMP9-satellite cell colocalization FC was analyzed by Mann-Whitney test following removal of one outlier from the OA group (ROUT method, Q = 0.01%). To assess whether MMP9 positivity was associated with exercise and/or satellite cell incarceration, a Fisher’s exact test was performed on satellite cell counts (pooled between groups) from pre- and post-exercise biopsies. Percent NCAM+ myofibres were assessed by Mann-Whitney test.

As a sensitivity analysis, data were log-transformed and reanalyzed without outlier removal. The results were consistent with the primary analysis. Data are expressed as mean ± standard deviation. As sex was unequally distributed between groups, a sensitivity analysis was conducted using ANCOVA with sex as a covariate to assess its potential effect on the absolute and normalized to body weight Biodex data. The results were consistent with the primary analysis.

## Results

### Participant Characteristics

See Table 5 for a detailed description of participant characteristics. Age, BMI, body fat percentage and TLM were similar between groups. No significant differences in walking, total aerobic or total resistance activity (minutes/week) were demonstrated between groups, however, the OA group reported participating in a higher number of minutes of lower-body resistance activity per week (*p* = 0.030). One participant in the OA group reported 150-minutes of lower-body exercise per week but was still included in analysis, despite exclusion criteria. No differences in grip strength (percentiles or absolute) or VO_2peak_ (percentiles or normalized to body weight or LBM) were demonstrated between groups. The OA group had significantly higher WOMAC scores (pain, *p* < 0.001; stiffness, *p* < 0.001; physical function, *p* < 0.001; total, *p* < 0.001) than the control group. All participants in the OA group had a total WOMAC score ≥ 40 except for two who had scores of 38 and 39. The number of medications taken regularly did not differ between groups.

**Table 5:** Participant Characteristics. Data are shown as mean ± standard deviation. * denotes a significant difference between groups (*p* < 0.05). Aerobic activity is a combination of low, moderate and high-intensity activity. One control participant’s activity data was not included in analysis as days of activity were reported rather than minutes. BMI, body mass index; BW, bodyweight; LBM, lean body mass; VO_2peak_, maximal volume of oxygen consumption; WOMAC, Western Ontario and McMaster Universities Osteoarthritis Index.

|  | Control (n = 16) | OA (n = 20) | p-value |
| --- | --- | --- | --- |
| Age | 65.7 ± 9.1 | 67.1 ± 7.2 | 0.628 |
| Sex Assigned at Birth, #female, #male | 8, 8 | 14, 6 |  |
| Height, cm | 170.3 ± 9.5 | 167.9 ± 9.5 | 0.465 |
| Weight, kg | 78.9 ± 12.0 | 73.2 ± 15.2 | 0.217 |
| BMI, kg/m<sup>2</sup> | 27.1 ± 3.0 | 25.8 ± 3.7 | 0.228 |
| Bodyfat Percentage | 36.2 ± 5.5 | 34.9 ± 7.2 | 0.548 |
| Thigh Lean Mass (kg) | 10.3 ± 1.4 | 9.6 ± 2.1 | 0.225 |
| Aerobic Activity (min/week) | 154.5 ± 195.6 | 167.1 ± 258.6 | 0.599 |
| Total Resistance Activity (min/week) | 27.3 ± 44.6 | 78.4 ± 89.9 | 0.094 |
| Lower Body Resistance Activity (min/week)* | 10.7 ± 22.3 | 38.4 ± 43.2 | 0.030 |
| Walking (min/week) | 136.0 ± 108.5 | 218.5 ± 586.4 | 0.195 |
| Grip Strength, kg | 70.1 ± 19.0 | 59.8 ± 15.6 | 0.090 |
| Grip Strength, percentile | 71.6 ± 23.7 | 57.9 ± 28.4 | 0.136 |
| $VO_{2peak}$, mL/min/kgBW | 26.9 ± 4.5 | 24.3 ± 6.7 | 0.200 |
| $VO_{2peak}$, mL/min/kgLBM | 43.9 ± 5.6 | 40.1 ± 10.3 | 0.162 |
| $VO_{2peak}$, percentile | 92.2 ± 11.4 | 78.6 ± 26.9 | 0.192 |
| WOMAC Pain* | 0.4 ± 1.0 | 10.9 ± 2.5 | < 0.001 |
| WOMAC Stiffness* | 1.0 ± 1.0 | 4.6 ± 1.4 | < 0.001 |
| WOMAC Physical Function* | 1.5 ± 2.5 | 35.9 ± 7.3 | < 0.001 |
| WOMAC Total* | 2.9 ± 4.2 | 51.3 ± 9.6 | < 0.001 |
| Number of Medications Taken Regularly | 1.4 ± 1.4 | 1.8 ± 1.4 | 0.336 |

### Knee Extensor Performance and Muscle Quality was Reduced in Participants with Moderate-to-Severe Knee OA

See Table 6 for knee extensor performance scores. Knee extensor performance was assessed on a Biodex dynamometer. The control group had greater absolute isometric peak force (*p* = 0.002) as well as force normalized to body weight (*p* = 0.006). Similarly, the control group has greater absolute isokinetic concentric peak torque (*p* < 0.001) as well as normalized to body weight (*p* = 0.008). To measure muscle quality (specific force), isometric and isokinetic scores were normalized to TLM. The control group had greater isometric (*p* = 0.002) and isokinetic (*p* = 0.002) muscle quality. RTD was not different between groups for absolute scores (*p* = 0.081), scores normalized to body weight (*p* = 0.120) or TLM (*p* = 0.103).

**Table 6:** Knee Extensor Performance and Muscle Quality from Volunteers with and without Moderate-to-Severe Knee OA. Data are shown as mean ± standard deviation. * denotes a significant difference between groups (*p* < 0.05). Peak isometric contractions were performed at a knee angle of 60° and peak isokinetic concentric contractions were performed at 60°/s. The following data were excluded from analysis due to measurement error or missing data: isometric data (*n* = 1 OA), isokinetic concentric data (*n* = 2 control; *n* = 2 OA), RTD (*n* = 3 control; *n* = 4 OA). BW, bodyweight; RTD, rate of torque development; TLM, thigh lean mass.

|  | Control | OA | *p*-value |
| --- | --- | --- | --- |
| Isometric (Nm)* | 143.1 ± 27.7 | 105.2 ± 37.7 | 0.002 |
| Isometric (Nm/kgBW)* | 1.8 ± 0.3 | 1.5 ± 0.4 | 0.006 |
| Isometric (Nm/kgTLM)* | 21.6 ± 3.8 | 17.4 ± 3.8 | 0.002 |
| Isokinetic Concentric (Nm)* | 118.3 ± 32.2 | 78.5 ± 25.9 | < 0.001 |
| Isokinetic Concentric (Nm/kgBW)* | 1.5 ± 0.4 | 1.1 ± 0.4 | 0.008 |
| Isokinetic Concentric (Nm/kgTLM)* | 17.6 ± 3.3 | 13.4 ± 3.8 | 0.002 |
| RTD (Nm/ms) | 1.0 ± 0.4 | 0.7 ± 0.4 | 0.081 |
| RTD (Nm/s/kgBW) | 12.1 ± 3.5 | 9.9 ± 3.8 | 0.120 |
| RTD (Nm/s/kgTLM) | 140.8 ± 39.9 | 116.5 ± 36.4 | 0.103 |

### Volunteers with Moderate-to-Severe Knee OA had Significantly more Fibrillar Muscle Collagen Area Fraction than Controls

Fibrillar collagen in skeletal muscle was assessed by Picrosirius red and expressed as a percentage of the total area analyzed (Figure 2). Participants with knee OA had significantly more fibrillar collagen content (12.2 ± 4.8%) than controls (8.4 ± 2.9%, *p* = 0.006).

**Figure 2:**
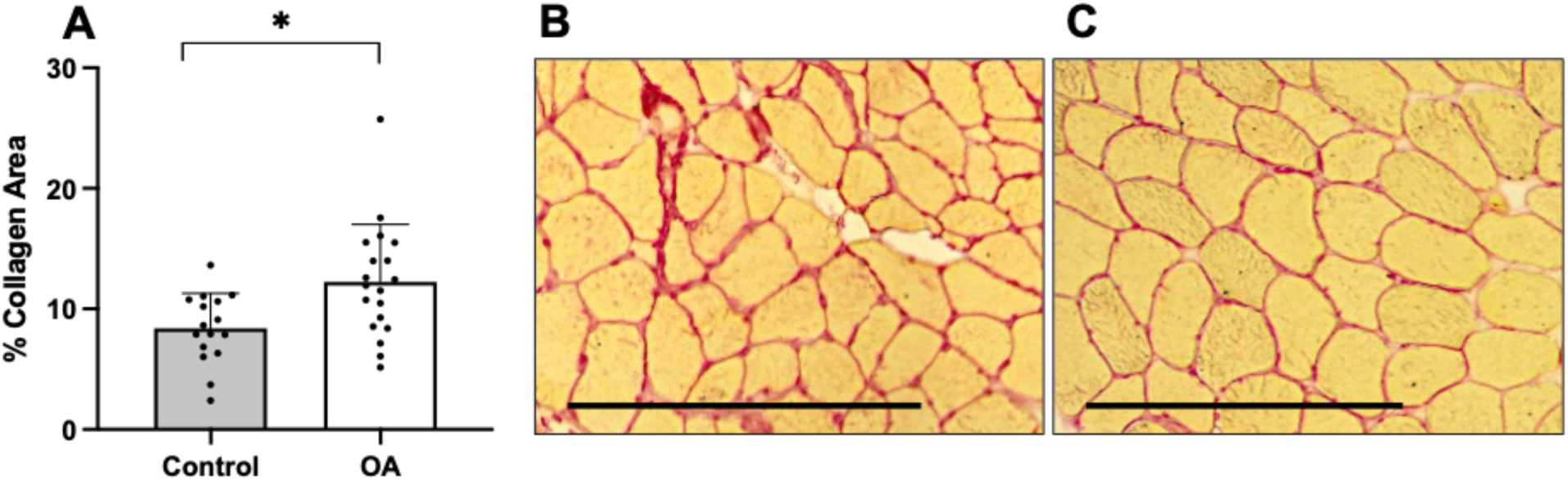
Fibrillar Muscle Collagen in Participants with and without Moderate-to-Severe Knee OA. *A)* Fibrillar muscle collagen is presented as mean percentage of total muscle area measured; volunteers with knee OA (*n* = 19) had significantly more fibrillar muscle collagen content than controls (*n* = 16, *p* = 0.006). Representative images of *B)* OA and *C)* control muscle where collagen is stained red. Error bar shows standard deviation. * denotes a significant difference between groups (*p* < 0.05). Scale bar = 375 μm. OA, osteoarthritis.

### Myofibre Morphology, Fibre Type Distribution and Capillarization in Participants with and without Moderate-to-Severe Knee OA

See Table 7 for a summary of myofibre morphology and fibre type distribution and Figure 3 for representative images. A main group effect was demonstrated in which participants with knee OA had significantly smaller muscle fibres (type I, 4728 ± 1755 μm^2^; type II, 3834 ± 2022 μm^2^) than control participants (type I, 6416 ± 1546 μm2; type II 5406 ± 2259 μm^2^, *p* < 0.001). A main effect of fibre type was also demonstrated in which type II muscle fibres were smaller than type I fibres (*p* = 0.011). No group differences in fibre distribution as expressed as percentage of fibres analyzed or percent CSA were demonstrated, however, both groups had a greater proportion of their muscle CSA comprised of type I myofibres (control, 55.3 ± 9.6 %; OA, 57.9 ± 8.3 %, *p* < 0.001). A main effect of group was demonstrated in which the volunteers with knee OA had a greater SFI than the control group (*p* = 0.039), indicating larger variability in fibre shape (Figure 3B). Capillaries per fibre were not significantly different between groups (control, 1.2 ± 0.3; OA, 1.1 ± 0.3, *p* = 0.276) (Figure 3C).

**Figure 3:**
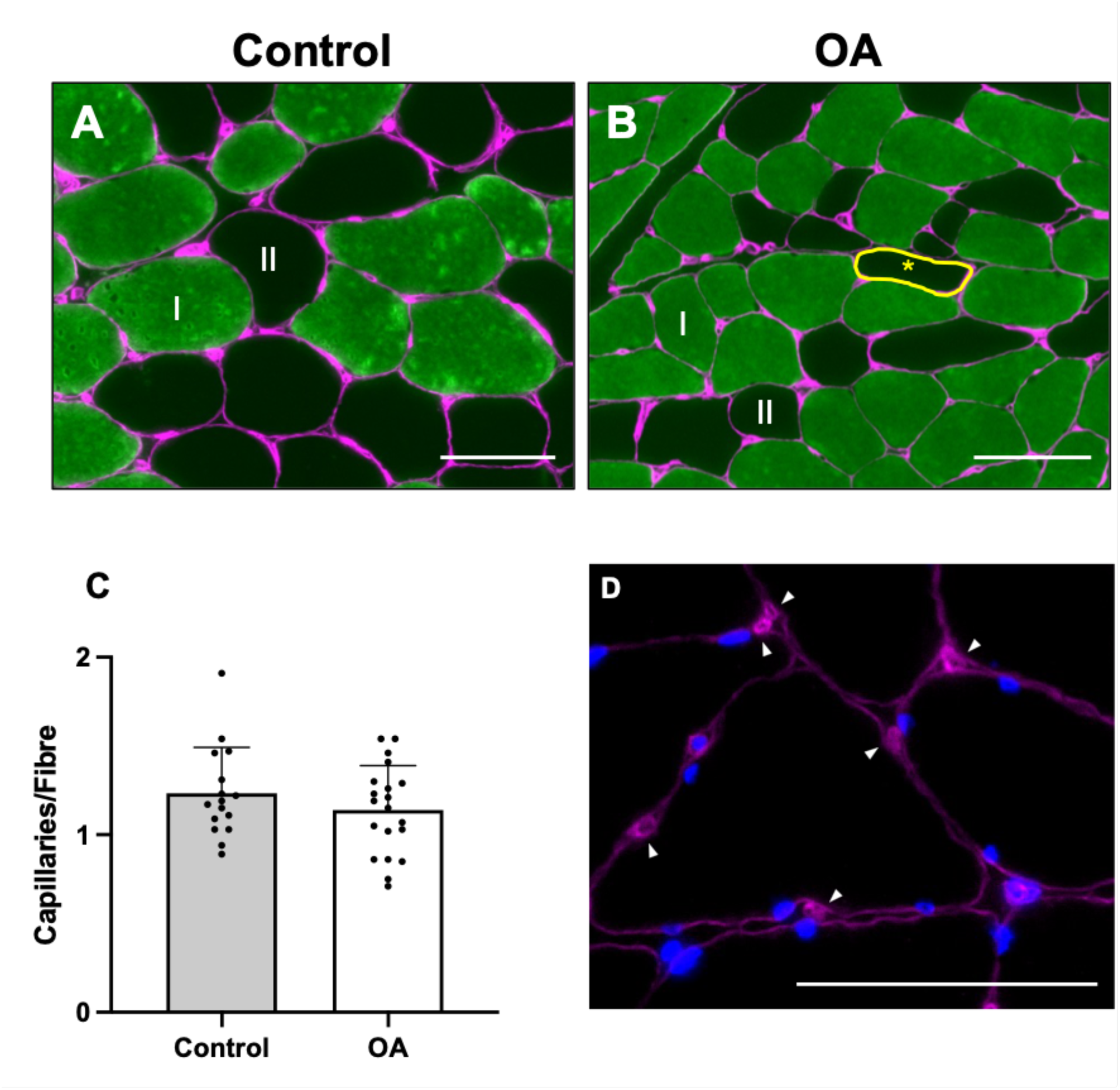
Myofibre Morphology and Capillaries in Participants with and without Moderate-to-Severe Knee OA. Representative images of muscle fibre-typing, shape factor index and capillary analyses. Green fibres are positive for MHC (slow fibres) (DSHB, A4.951) and are classified as type I fibres (I), whereas black fibres are MHC (slow fibres)-negative and are classified as type II fibres (II). *A)* Representative control muscle sample, type I myofibres (green), type II myofibres (black), laminin (magenta). *B)* Representative OA muscle sample, type I myofibres (green), type II myofibres (black), laminin (magenta). Yellow circled and asterisked fibre highlights a myofibre with a higher shape factor index (irregular fibre shape). *C)* No significant groups differences in capillaries per fibre were demonstrated (*n* = 16 control; *n* = 20 OA). *D)* Representative muscle capillarization image, laminin (magenta) + DAPI (blue). White arrowheads indicate presence of a capillary. Error bar shows standard deviation. Scale bar = 100 μm. OA, osteoarthritis.

**Table 7:** Myofibre Morphology and Fibre Type Distribution in Participants with and without Moderate-to-Severe Knee OA. Data are shown as mean ± standard deviation. Data presented below are complete (*n* = 16 control; *n* = 20 OA), except for one OA value excluded from the myofibre CSA analysis following identification as an outlier. # denotes a main effect of fibre type; § denotes a main effect of group (*p* < 0.05). CSA, cross-sectional area; OA, osteoarthritis.

|  | **Fibre Type** | **Control** | **OA** |
| --- | --- | --- | --- |
| **Myofibre CSA ($\mu\text{m}^2$)<sup>§</sup>** | I | 6416 ± 1546 | 4728 ± 1755 |
| | II | 5406 ± 2259 | 3834 ± 2022 |
| **Fibre Type Distribution (fibre %)** | I | 46.4 ± 16.0 | 51.4 ± 17.7 |
| | II | 53.6 ± 16.0 | 48.6 ± 17.7 |
| **Fibre Type Distribution (CSA %)<sup>#</sup>** | I | 55.3 ± 9.6 | 57.9 ± 8.3 |
| | II | 44.7 ± 9.6 | 42.1 ± 8.3 |
| **Shape Factor Index<sup>§</sup>** | I | 1.33 ± 0.05 | 1.37 ± 0.07 |
| | II | 1.36 ± 0.07 | 1.40 ± 0.10 |

### Measures of Knee Extensor Performance Negatively Correlated with Percent Fibrillar Muscle Collagen Content and SFI

Percent collagen area as measured by Picrosirius red was correlated with measures of knee extensor performance and reported in absolute measures to maintain consistency with previous reports (Noehren et al., 2018) (Figure 4). Peak isokinetic concentric torque was negatively correlated with percent collagen area (*p* = 0.006, *r* = -0.479) (Figure 4A) but not peak isometric force (*p* = 0.104, *r* = -0.283) (Figure 4B) or RTD (*p* = 0.089, *r* = -0.327) (Figure 4C).

**Figure 4:**
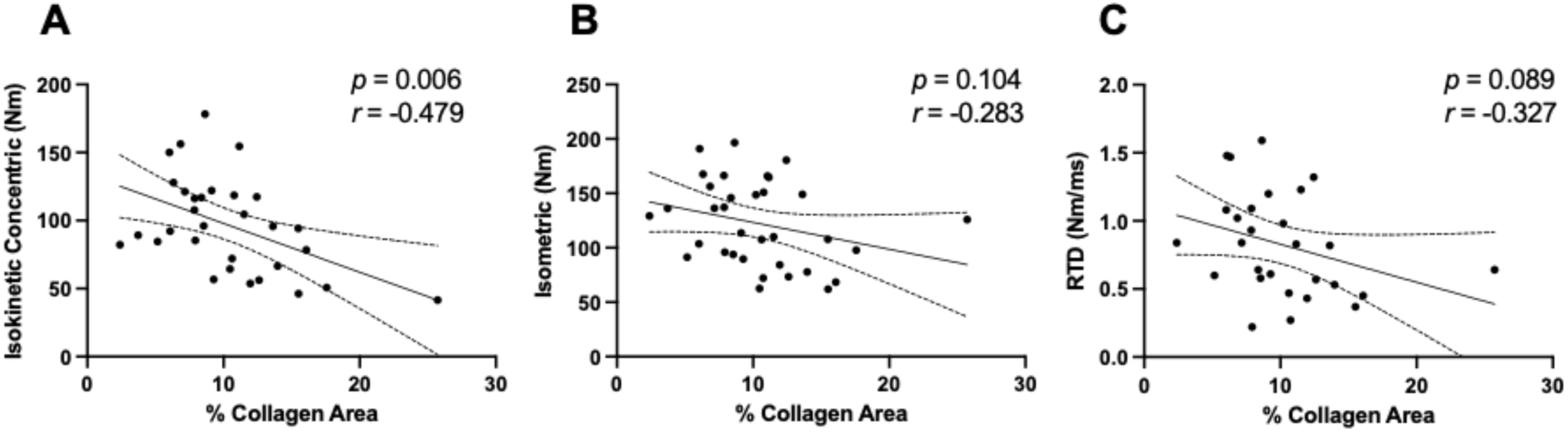
Knee Extensor Performance Correlations with Percent Muscle Collagen Area. *A)* Peak isokinetic concentric torque negatively correlated with percent muscle collagen area as measured by Picrosirius red. *B)* Peak isometric force with percent collagen area *C)* RTD with percent collagen area did not significantly correlate. Solid lines represent the line of best fit; dashed lines represent the 95% confidence interval. RTD, rate of torque development.

Measures of knee extensor performance were also correlated with SFI from both type I and type II myofibres (Figure 5). To maintain consistency with previous reports, data were reported in absolute terms as well as normalized to TLM (Soendenbroe et al., 2024). Type I and type II myofibre SFI negatively correlated with absolute isometric force (type I, *p* = 0.011, *r* = -0.432; type II, *p* = 0.005, *r* = -0.475) (Figure 5A) and isometric force relative to TLM (type I, *p* = 0.018, *r* = -0.405; type II, *p* = 0.002, *r* = -0.512) (Figure 5B). Type I but not type II myofibre SFI negatively correlated with absolute isokinetic peak torque (type I, *p* = 0.012, *r* = -0.444; type II, *p* = 0.153, *r* = -0.263) (Figure 5C) as well as isokinetic peak torque normalized to TLM (type I, *p* = 0.034, *r* = -0.382; type II, *p* = 0.285, *r* = -0.198) (Figure 5D). Similarly, RTD negatively correlated with type I myofibre SFI (*p* = 0.030, *r* = -0.412) but not type II myofibre SFI (*p* = 0.092, *r* = -0.324) (Figure 5E).

**Figure 5:**
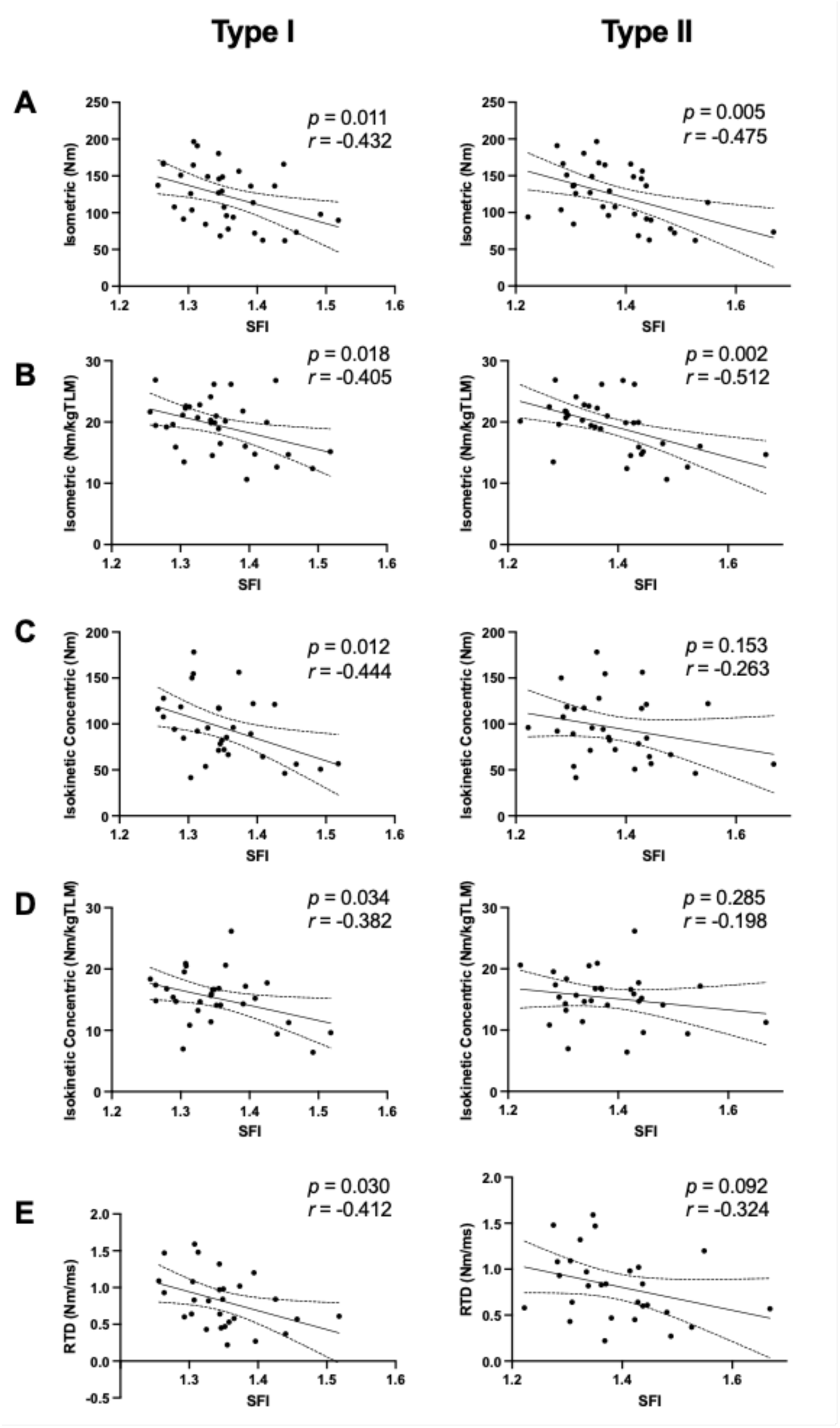
Knee Extensor Performance Correlations with SFI by Fibre Type. Peak absolute *A)* and specific *B)* isometric force negatively correlated with SFI from both type I and type II myofibres. *C)* Type I but not type II myofibre SFI negatively correlated with peak isokinetic concentric torque. *D)* Type I but not type II myofibre SFI negatively correlated with peak isokinetic concentric specific force. *E)* Type I but not type II myofibre SFI negatively correlated with RTD. Solid lines represent the line of best fit; dashed lines represent the 95% confidence interval. RTD, rate of torque development; TLM, thigh lean mass.

### *MMP2, MMP9, MMP14, TIMP1* and *TIMP2* Gene Expression Increased 72h after Acute Resistance Exercise in the Muscles from Volunteers with and without Moderate-to-Severe Knee OA

*MMP2, MMP9, MMP14, TIMP1* and *TIMP2* gene expression were quantified by qPCR, and results are reported as the FD from the average control-0h value (Figure 6; Figure 7). All genes demonstrated a main effect of time, *MMP2* (control-0h, 1.0 ± 0.3; OA-0h, 1.1 ± 0.4; control-72h, 2.3 ± 1.3; OA-72h, 1.9 ± 0.9 FD; *p* < 0.001) (Figure 6A), *MMP9* (control-0h, 1.5 ± 1.5; OA-0h, 0.8 ± 0.5; control-72h, 12.4 ± 15.5; OA-72h, 12.8 ± 17.0 FD; *p* = 0.001) (Figure 6B), *MMP14* (control-0h, 1.0 ± 0.4; OA-0h, 1.2 ± 0.6; control-72h, 3.8 ± 3.5; OA-72h, 3.9 ± 3.4 FD; *p* < 0.001) (Figure 6C), *TIMP1* (control-0h, 1.1 ± 0.5; OA-0h, 1.2 ± 0.4; control-72h, 8.3 ± 7.4; OA-72h, 5.8 ± 4.1 FD; *p* < 0.001) (Figure 7A), and *TIMP2* (control-0h, 1.0 ± 0.3; OA-0h, 1.2 ± 0.6; control-72h, 2.9 ± 2.5; OA-72h, 2.6 ± 2.0 FD; *p* = 0.002) (Figure 7B). There were no significant group effects.

**Figure 6:**
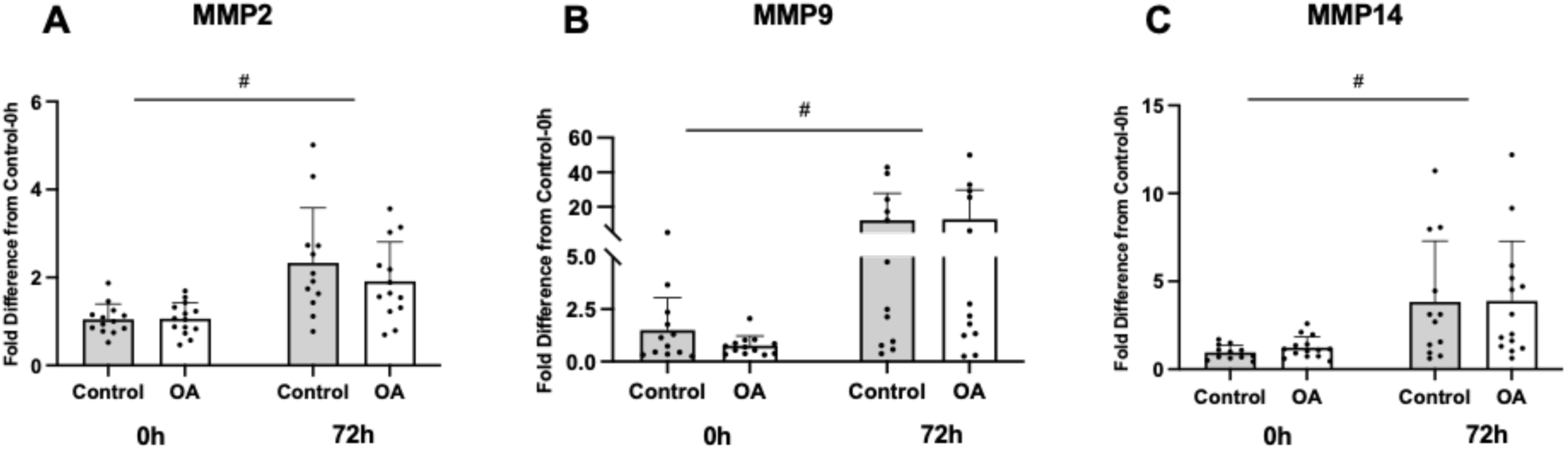
MMP Gene Expression 72h after Acute Resistance Exercise in Skeletal Muscle from Volunteers with and without Moderate-to-Severe Knee OA. *A) MMP2* (*n* = 13 control-0h; *n* = 12 control-72h; *n* = 14 OA-0h; *n* = 13 OA-72h; *p* < 0.001)*, B) MMP9* (*n* = 12 control-0h; *n* = 12 control-72h; *n* = 13 OA-0h; *n* = 12 OA-72h; *p* = 0.001), and *C) MMP14* (*n* = 12 control-0h; *n* = 12 control-72h; *n* = 14 OA-0h; *n* = 14 OA-72h; *p* < 0.001) gene expression were elevated 72h after acute resistance exercise irrespective of group. Data were normalized against the average control-0h value. Error bar shows standard deviation. # denotes a main effect of time (*p* < 0.05). 0h, pre-exercise data; 72h, 72 hours post-exercise data; MMP, matrix metalloproteinase; MMP2, matrix metalloproteinase-2; MMP9, matrix metalloproteinase-9; MMP14, matrix metalloproteinase-14; OA, osteoarthritis.

**Figure 7:**
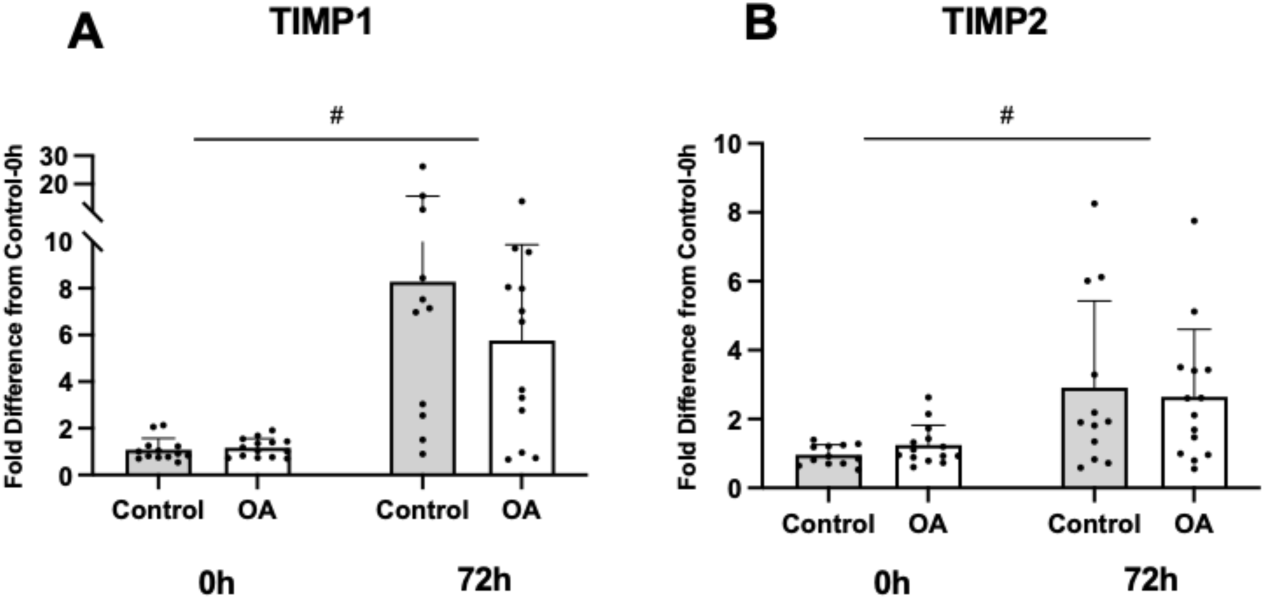
TIMP Gene Expression 72h after Acute Resistance Exercise in Skeletal Muscle from Volunteers with and without Moderate-to-Severe Knee OA. *A) TIMP1* (*n* = 13 control-0h; *n* = 11 control-72h; *n* = 14 OA-0h; *n* = 13 OA-72h; *p* < 0.001) and *B) TIMP2* (*n* = 12 control-0h; *n* = 12 control-72h; *n* = 14 OA-0h; *n* = 14 OA-72h; *p* = 0.002) gene expression were elevated 72h after acute resistance exercise irrespective of group. Data were normalized against the average control-0h value. Error bar shows standard deviation. # denotes a main effect of time (*p* < 0.05). 0h, pre-exercise data; 72h, 72 hours post-exercise data; TIMP, tissue inhibitor of metalloproteinase; TIMP1, tissue inhibitor of metalloproteinase-1; TIMP2, tissue inhibitor of metalloproteinase-2; OA, osteoarthritis.

### MMP2 and MMP9 Protein Response to Acute Resistance Exercise were Altered in the Muscle ECM from Volunteers with Moderate-to-Severe Knee OA

MMP2 and MMP9 protein were analyzed through immunofluorescent imaging and expressed as a percentage of the total area analyzed (Figure 8; Figure 9). For MMP2, the mixed effects model demonstrated a main effect of time (*p* = 0.002) and group (*p* = 0.010) as well as a group by time interaction (*p* = 0.041). In the control group, compared to baseline (4.1 ± 2.3%), MMP2 protein content was elevated 72h after exercise (6.3 ± 3.3%, *p* = 0.001). Control-72h MMP2 protein content was also greater than OA-72h MMP2 protein content (3.3 ± 1.8%, *p* = 0.001). Baseline OA MMP2 protein content (2.9 ± 1.7%) did not differ from control baseline or OA-72h content (Figure 8A). Similarly, MMP9 protein content demonstrated a main effect of time (*p* = 0.001) and group (*p* < 0.001) as well as a group by time interaction (*p* = 0.004). Control-72h MMP9 content (1.5 ± 1.2%) was significantly greater than baseline control content (0.6 ± 0.3%, *p* < 0.001) and OA-72h content (0.4 ± 0.2%, *p* < 0.001), whereas baseline OA content (0.3 ± 0.2%) did not significantly differ from any group (Figure 9A). MMP9 colocalization to capillaries was evident visually in both the 0h and 72h post-exercise time points (Figure 9F, Figure 9G).

**Figure 8:**
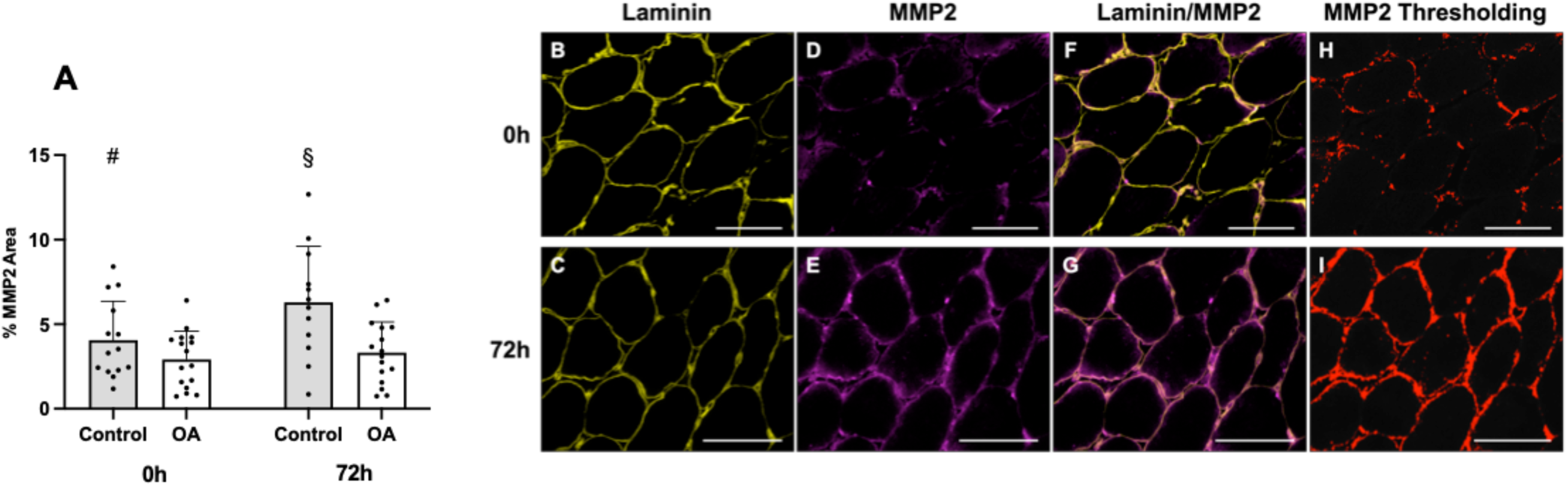
MMP2 Protein Fraction in Skeletal Muscle from Volunteers with and without Moderate-to-Severe Knee OA. *A)* Quantification of MMP2 protein is presented as the mean percentage of total muscle area analyzed (*n* = 14 control-0h; *n* = 12 control-72h; *n* = 16 OA-0h; *n* = 16 OA-72h). In the control group, compared to baseline (0h), MMP2 area fraction was elevated 72h after acute resistance exercise (*p* = 0.001). 72h after resistance exercise, MMP2 area fraction in the control group was greater than the OA group (*p* = 0.001). Error bar shows standard deviation. # denotes significant difference from the control-72h time point; § denotes significant difference from the OA-72h time point (*p* < 0.05). *B-G)* Representative images of Laminin and MMP2 staining in skeletal muscle, before (0h) and 72h after acute resistance exercise. *B/C)* Laminin (yellow). *D/E)* MMP2 (magenta). *F/G)* Laminin (yellow) + MMP2 (magenta). *H/I)* Representation of thresholding analysis used to calculate percent MMP2 area. Analysis was performed with ImageJ software with thresholds kept constant between all samples. Red area represents MMP2-positive staining after thresholding and corresponds to immunofluorescent staining in figures D/E. Scale bar = 100 μm. 0h, pre-exercise time point; 72h, 72 hours post-exercise time point; MMP2, matrix metalloproteinase-2; OA, osteoarthritis.

**Figure 9:**
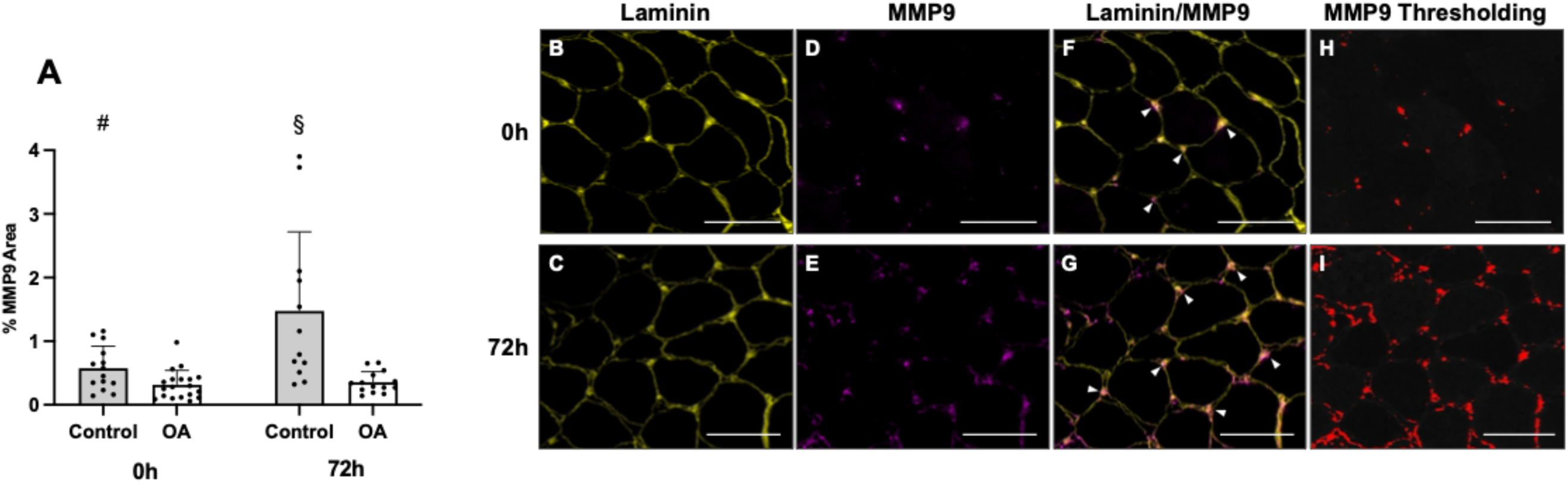
MMP9 Protein Fraction in Skeletal Muscle from Volunteers with and without Moderate-to-Severe Knee OA. *A)* Quantification of MMP9 protein is presented as the mean percentage of total muscle area analyzed (*n* = 14 control-0h; *n* = 12 control-72h; *n* = 19 OA-0h; *n* = 14 OA-72h). In the control group, compared to baseline (0h), MMP9 area fraction was elevated 72h after acute resistance exercise (*p* < 0.001). 72h after resistance exercise, MMP9 area fraction in the control group was greater than the OA group (*p* < 0.001). Error bar shows standard deviation. # denotes a significant difference from the control-72h time point; § denotes a significant difference from the OA-72h time point (*p* < 0.05). *B-G)* Representative images of Laminin and MMP9 staining in skeletal muscle, pre (0h) and 72h after acute resistance exercise. *B/C)* Laminin (yellow). *D/E)* MMP9 (magenta). *F/G)* Laminin (yellow) + MMP9 (magenta), white arrowheads show colocalization of MMP9 to capillaries. *H/I)* Representation of thresholding analysis used to calculate percent MMP9 area. Analysis was performed with ImageJ software with thresholds kept constant between all samples. Red area represents MMP9-positive staining after thresholding and corresponds to immunofluorescent staining in figures D/E. Scale bar = 100 μm. 0h, pre-exercise time point; 72h, 72 hours post-exercise time point; MMP9, matrix metalloproteinase-9; OA, osteoarthritis.

### MMP2 and MMP9 Enzyme Activity Potential Increased after Acute Resistance Exercise in Muscles from Volunteers with and without Moderate-to-Severe Knee OA

MMP2 and MMP9 enzyme activity potential were quantified by substrate zymography (Figure 10). For MMP2, the mixed effects model demonstrated a main effect of time (control, 1.5 ± 0.5 FC; OA, 1.9 ± 0.9 FC, *p* < 0.001) (Figure 10A). Similarly, the mixed effects model demonstrated a main effect of time for MMP9 (control, 2.0 ± 1.6; OA, 1.6 ± 0.7 FC, *p* = 0.001) (Figure 10B). No group effect was demonstrated for either enzyme.

**Figure 10:**
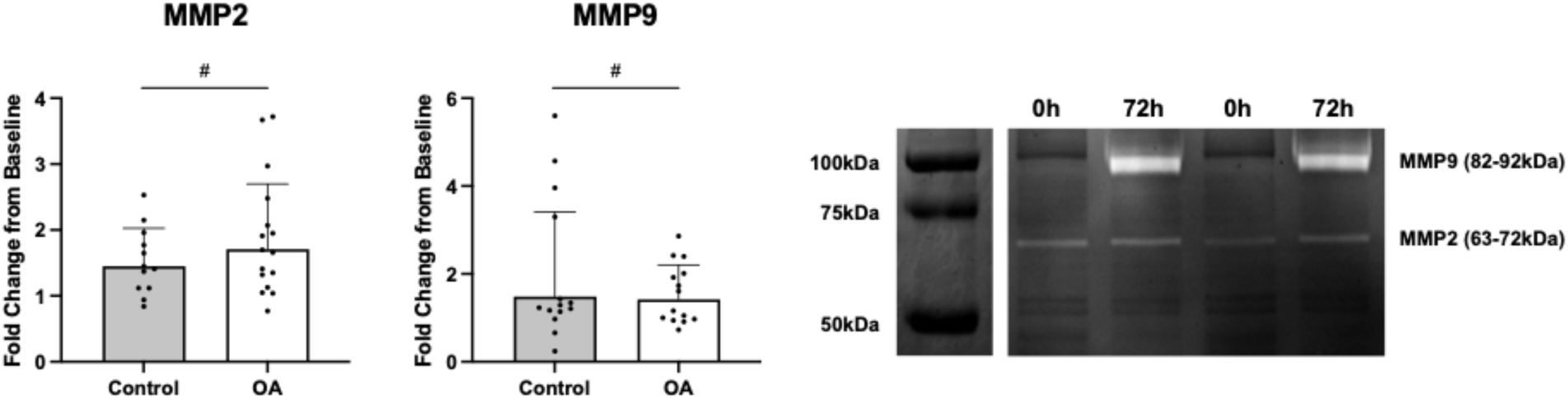
MMP2 and MMP9 Enzyme Activity Potential 72h After Acute Resistance Exercise in Skeletal Muscle from Volunteers with and without Moderate-to-Severe Knee OA. *A)* MMP2 (*n* = 12 control; *n* = 16 OA; *p* < 0.001) and *B)* MMP9 (*n* = 14 control; *n* = 14 OA; *p* = 0.001) enzyme activity potential were elevated 72h after acute resistance exercise in both groups. Error bar shows standard deviation. # denotes a main effect of time when baseline is set to 1 (*p* < 0.05). *C)* Representative zymogram; areas of brighter white represent areas of greater digestion and enzyme activity. MMP2, matrix metalloproteinase-2; MMP9, matrix metalloproteinase-9; OA, osteoarthritis.

### Volunteers with Moderate-to-Severe Knee OA had more PDGFRα+ Mesenchymal Cells and these Cells Positively Correlated with Fibrillar Muscle Collagen and Negatively Correlated with MMP9 Area Fraction

PDGFRα+ mesenchymal cells were quantified by immunofluorescent imaging and expressed as the number of cells present per mm^2^ of tissue (Figure 11). A main effect of group was observed, with volunteers in the OA group having more PDGFRα+ mesenchymal cells (0h, 11.0 ± 4.0; 72h, 10.9 ± 3.2 PDGFRα+/mm^2^) than controls (0h, 6.7 ± 1.4; 72h, 8.3 ± 3.5 PDGFRα+/mm^2^, *p* < 0.001) (Figure 11A). No significant effect of exercise on PDGFRα+ mesenchymal cells was demonstrated; therefore, PDGFRα+ mesenchymal cell counts were collapsed across time for linear regression analyses. PDGFRα+ mesenchymal cells per mm^2^ positively correlated with fibrillar muscle collagen content (*p* = 0.029, *r* = 0.381) (Figure 11B). PDGFRα+ mesenchymal cells negatively correlated with MMP9 percent area, both at the baseline (*p* = 0.009, *r* = -0.455) (Figure 11C) and 72h post-exercise (*p* = 0.004, *r* = -0.561) (Figure 11D) but did not significantly correlate with MMP2 percent area at either time point.

**Figure 11:**
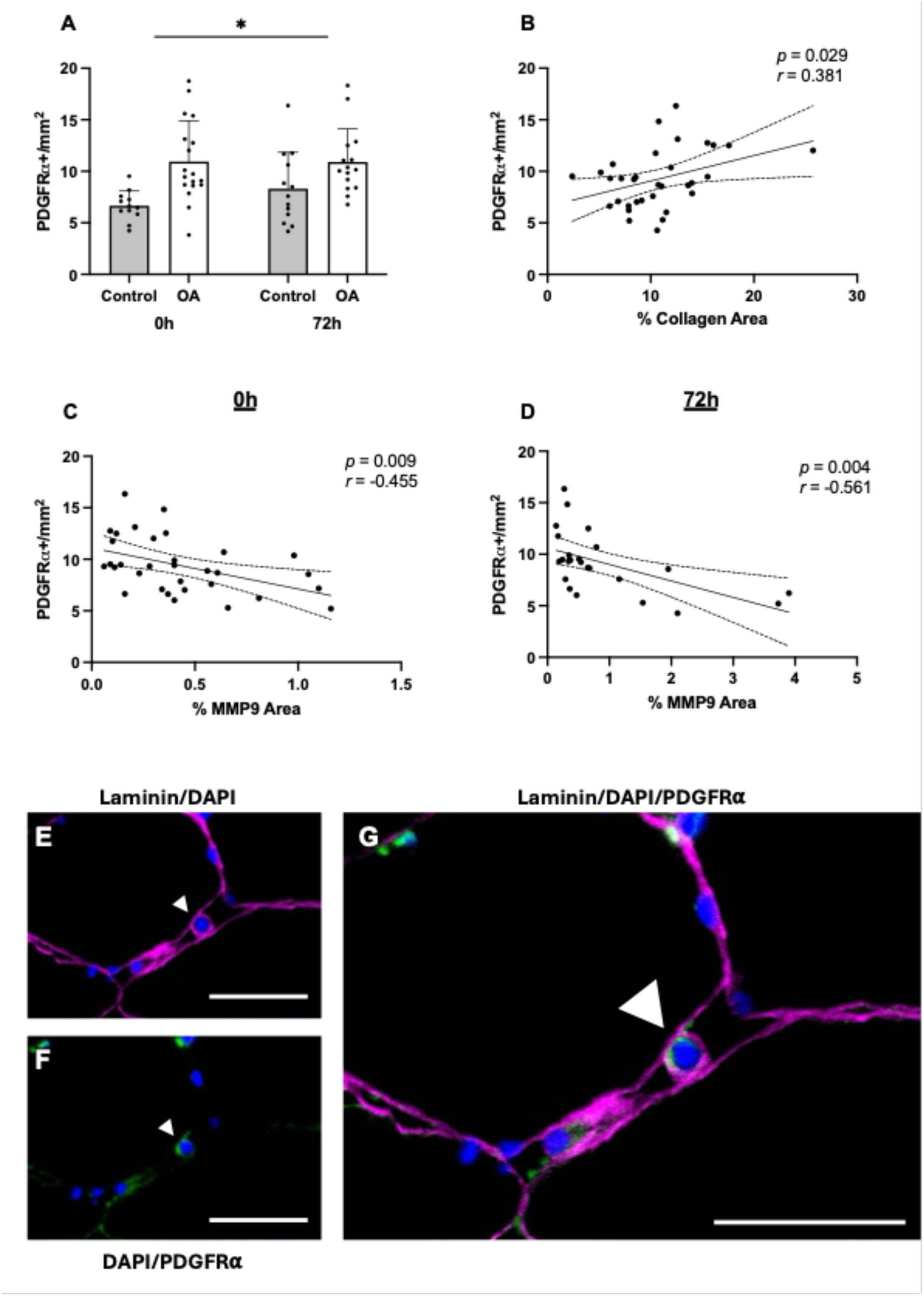
PDGFRα+ Mesenchymal Cells in Skeletal Muscle from Volunteers with and without Moderate-to-Severe Knee OA and Correlations with Fibrillar Muscle Collagen and MMP9 Area Fraction. *A)* A main group effect demonstrated that participants with knee OA had more PDGFRα+ cells per mm^2^ than controls (*n* = 12 control-0h; *n* = 13 control-72h; *n* = 18 OA-0h; *n* = 15 OA-72h; *p* < 0.001). No effect of exercise on PDGFRα+ cells was observed. Error bar shows standard deviation. * denotes a main group effect (*p* < 0.05). *B)* PDGFRα+ cells per mm^2^ positively correlated with fibrillar muscle collagen fraction (measured by Picrosirius red). PDGFRα+ cells per mm^2^ negatively correlated with MMP9 area fraction both at *C)* baseline (0h) and *D)* 72h post-exercise. Solid lines represent the line of best fit; dashed lines represent the 95% confidence interval. *E)* Laminin (magenta) + DAPI (blue), white arrowhead indicates PDGFRα+ mesenchymal cell. *F)* DAPI (blue) + PDGFRα (green), white arrowhead indicates PDGFRα+ mesenchymal cell. *G)* Laminin (magenta) + DAPI (blue) + PDGFRα (green), white arrowhead indicates PDGFRα+ mesenchymal cell. Scale bar = 50 μm. 0h, pre-exercise time point; 72h, 72 hours post-exercise time point; MMP9, matrix metalloproteinase-9; OA, osteoarthritis; PDGFRα, platelet-derived growth factor receptor-alpha.

### Volunteers with Moderate-to-Severe Knee OA had Less Satellite Cells

Satellite cells, as defined by colocalization of Pax7 and DAPI beneath the basal lamina of the myofibre (Nederveen et al., 2016), were quantified by immunofluorescent imaging (Table 8; Figure 12). Satellite cells were assessed before (0h) and 72h after acute resistance exercise and was expressed per type I or type II myofibre. A main group effect was observed with participants with moderate-to-severe knee OA having less satellite cells than controls (*p* = 0.006). A main effect of fibre type demonstrated that both groups had significantly fewer satellite cells associated with type II than type I myofibres (*p* < 0.001). No effect of exercise was demonstrated. Satellite cells with the lamina covering ≥ 90% of their perimeter were classified as “incarcerated” (Nederveen et al., 2020) (Figure 12). No significant difference in incarcerated satellite cells was demonstrated between group or fibre type (Table 8).

**Figure 12:**
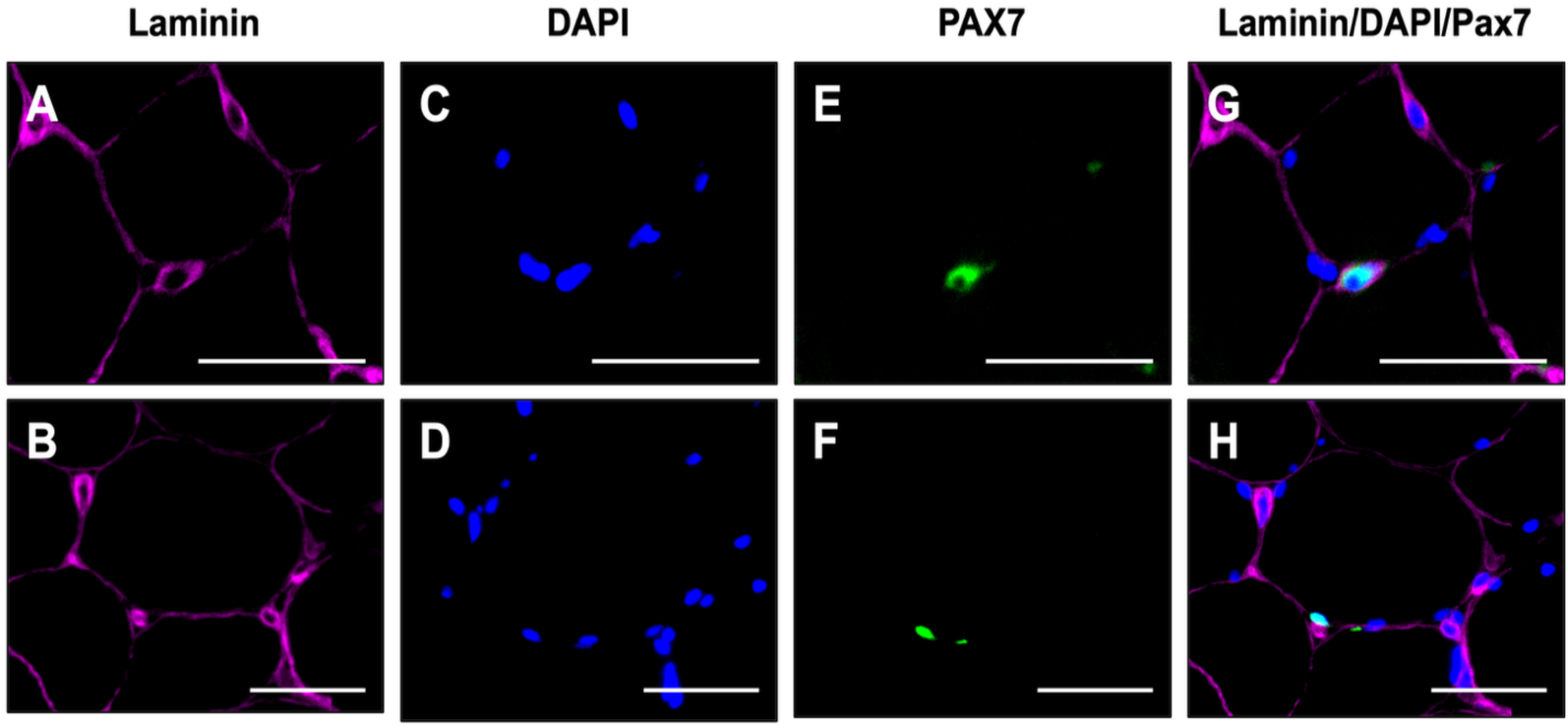
Representative Images of an Incarcerated and Non-incarcerated Satellite Cell. Satellite cells with lamina covering ≥ 90% of their perimeter were classified as incarcerated (Nederveen et al., 2020). *A/B)* Laminin (magenta). *C/D)* DAPI (blue). *E/F)* Pax7 (green). *G)* Representation of an incarcerated satellite cell. Laminin (magenta) + DAPI (blue) + Pax7 (green), DAPI + Pax7 overlay (cyan). *H)* Representation of a non-incarcerate satellite cell. Laminin (magenta) + DAPI (blue) + Pax7 (green), DAPI + Pax7 overlay (cyan). Scale bar = 50 μm.

**Table 8:**
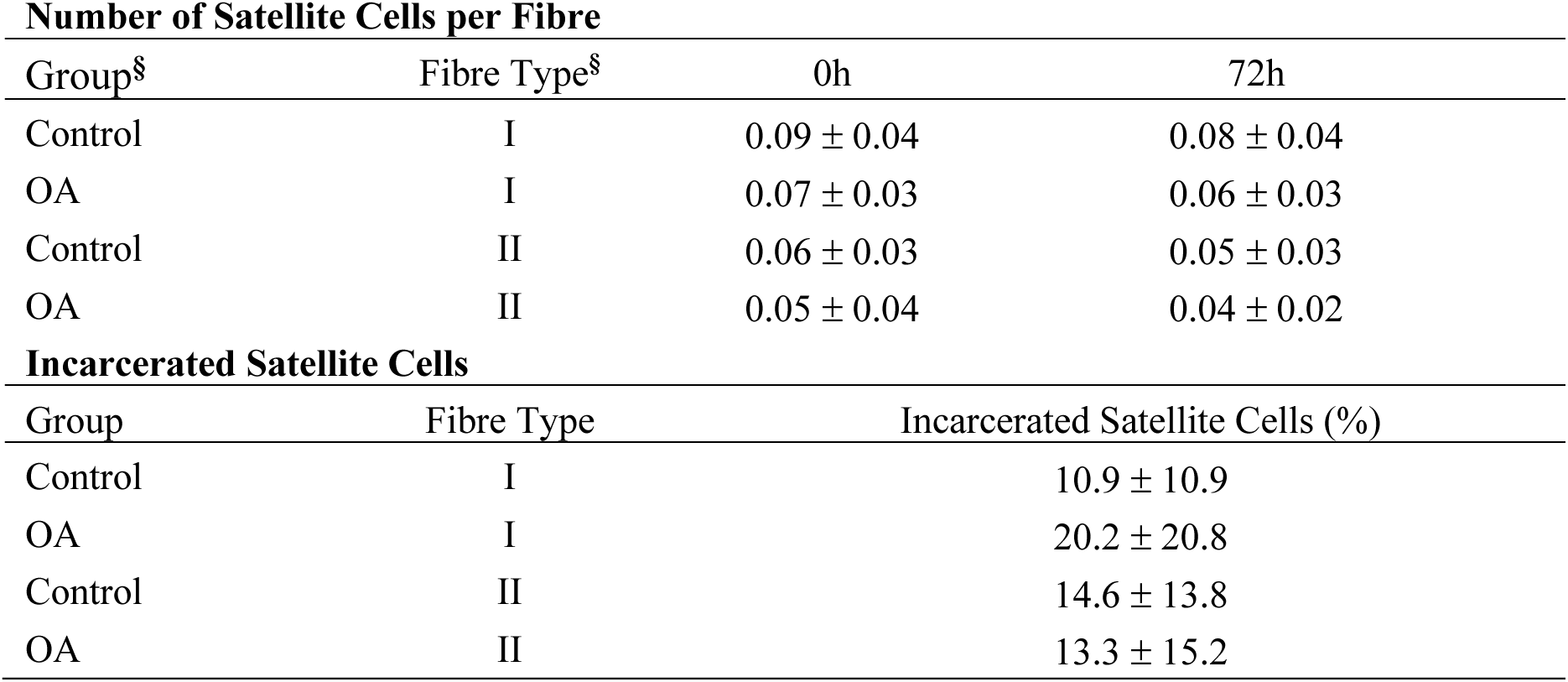
Satellite Cells Before and 72h after Acute Resistance Exercise and Incarcerated Satellite Cells in Volunteers with and without Moderate-to-Severe Knee OA. Data are presented as mean ± standard deviation. For analysis of number of satellite cells per fibre, sample sizes are as follows for both type I and type II fibres: *n* = 16 control-0h; *n* = 12 control-72h; *n* = 19 OA-0h; *n* = 17 OA-72h. For incarcerated satellite cell analysis, sample sizes are as following: *n* = 15 control (type I); *n* = 15 control (type II); *n* = 19 OA (type I); *n* = 17 OA (type II). § denotes a main effect (*p* < 0.05). 0h, pre-exercise time point; 72h, 72 hours post-exercise time point; OA, osteoarthritis.

### Individual Fold Change in MMP9+ Satellite Cells was Altered in Muscles from Volunteers with Moderate-to-Severe Knee OA 72h after Acute Resistance Exercise

Satellite cell-MMP9 colocalization was determined by immunofluorescence imaging (Figure 1), and MMP9+ satellite cells were quantified as a percentage of total satellite cells (Figure 13). The mixed effects model demonstrated no effect of group or time (Figure 13A), however, when individual response to acute resistance exercise was assessed (72h FC), the control group showed a greater response to exercise (1.5 ± 1.1 FC) than the OA group (0.8 ± 0.7 FC, *p* = 0.048) (Figure 13B).

**Figure 13:**
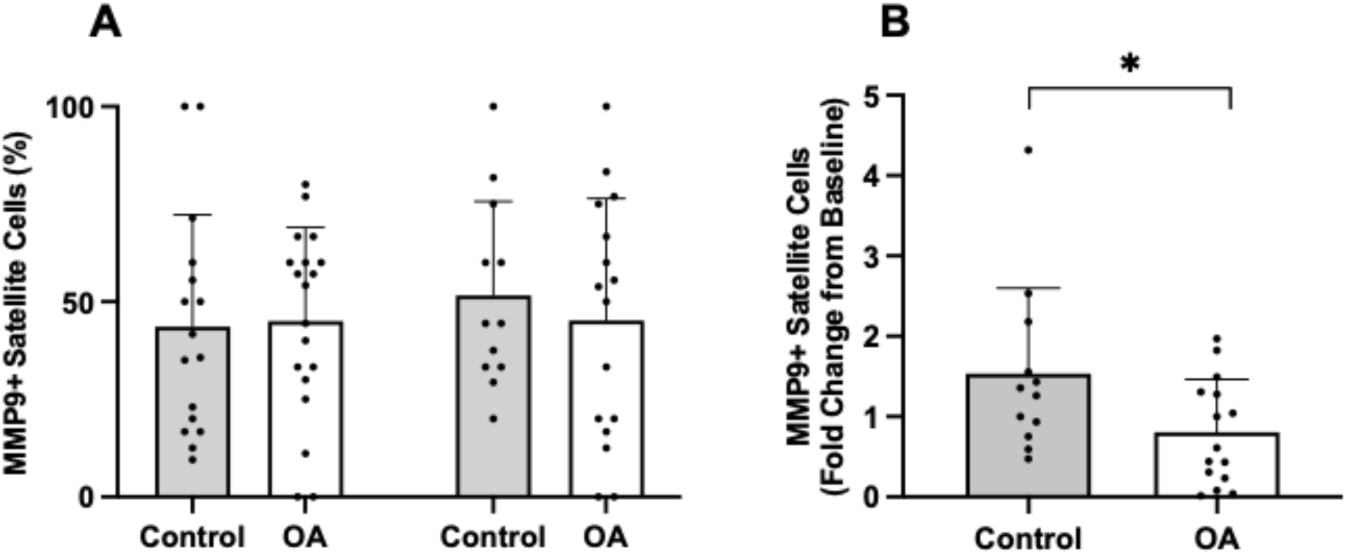
Satellite Cell-MMP9 Colocalization Before and 72h after Acute Resistance Exercise in Muscles from Volunteers with and without Moderate-to-Severe Knee OA. Satellite cell-MMP9 colocalization (MMP9+ satellite cells) was assessed by co-staining of Pax7 and MMP9 antibodies. *(A)* Data are expressed as the percentage of total satellite cells assessed to be MMP9+. Percent MMP9+ satellite cell content did not differ between group or time point (*n* = 16 control-0h; *n* = 12 control-72h; *n* = 19 OA-0h; *n* = 16 OA-72h). *(B)* The individual fold change in MMP9+ satellite cell content from pre-to 72h post-exercise was greater in the control group than the OA group (*n* = 12 control; *n* = 15 OA; *p* = 0.048). Error bar shows standard deviation. * denotes a significant difference between groups (*p* < 0.05).

### Satellite Cells were more Likely to be MMP9+ than MMP9-72h after Acute Resistance Exercise and Incarcerated Satellite Cells were more Likely to be MMP9+ than Non-incarcerated Cells

A Fisher’s exact test was run on satellite cell counts pooled between groups from pre-exercise (*n* = 378) and post-exercise (*n* = 232) biopsies to determine if there was an association between exercise and MMP9 positivity. A higher proportion of satellite cells colocalized to MMP9 72h after exercise (50.0%) compared to before exercise (41.3%, *p* = 0.036) (odds ratio [OR] = 1.423; 95% confidence interval [CI] [1.030, 1.970]). As a follow-up, a Fisher’s exact test was run on satellite cell counts pooled between groups from pre-exercise (*n* = 314 non-incarcerated; *n* = 64 incarcerated) and post-exercise (*n* = 202 non-incarcerated; *n* = 30 incarcerated) biopsies to determine if there was an association between incarceration and MMP9 positivity. A higher proportion of incarcerated satellite cells colocalized to MMP9 (70.0%) compared to non-incarcerated (47.0%, *p* = 0.030) 72h after acute resistance exercise (OR = 2.628; 95% CI [1.166, 6.247]). The proportion of satellite cells colocalizing to MMP9 from pre-exercise biopsies was not significantly different between incarcerated (51.6%) and non-incarcerated cells (31.2%, *p* = 0.072; OR = 1.653; 95% CI [0.967, 2.843]).

### NCAM+ Fibres were Rare with No Differences between Groups

NCAM+ myofibres were quantified by immunofluorescence and expressed as a percent of the total fibres analyzed per participant. Of the 10 control samples analyzed, three had NCAM+ staining and of the 17 OA samples analyzed, one had NCAM+ staining. No group differences were demonstrated in the percentage of myofibres that stained positive for NCAM (control, 0.5 ± 1.2%; OA, 0.08 ± 0.3%, *p* = 0.234) (Figure 14).

**Figure 14:**
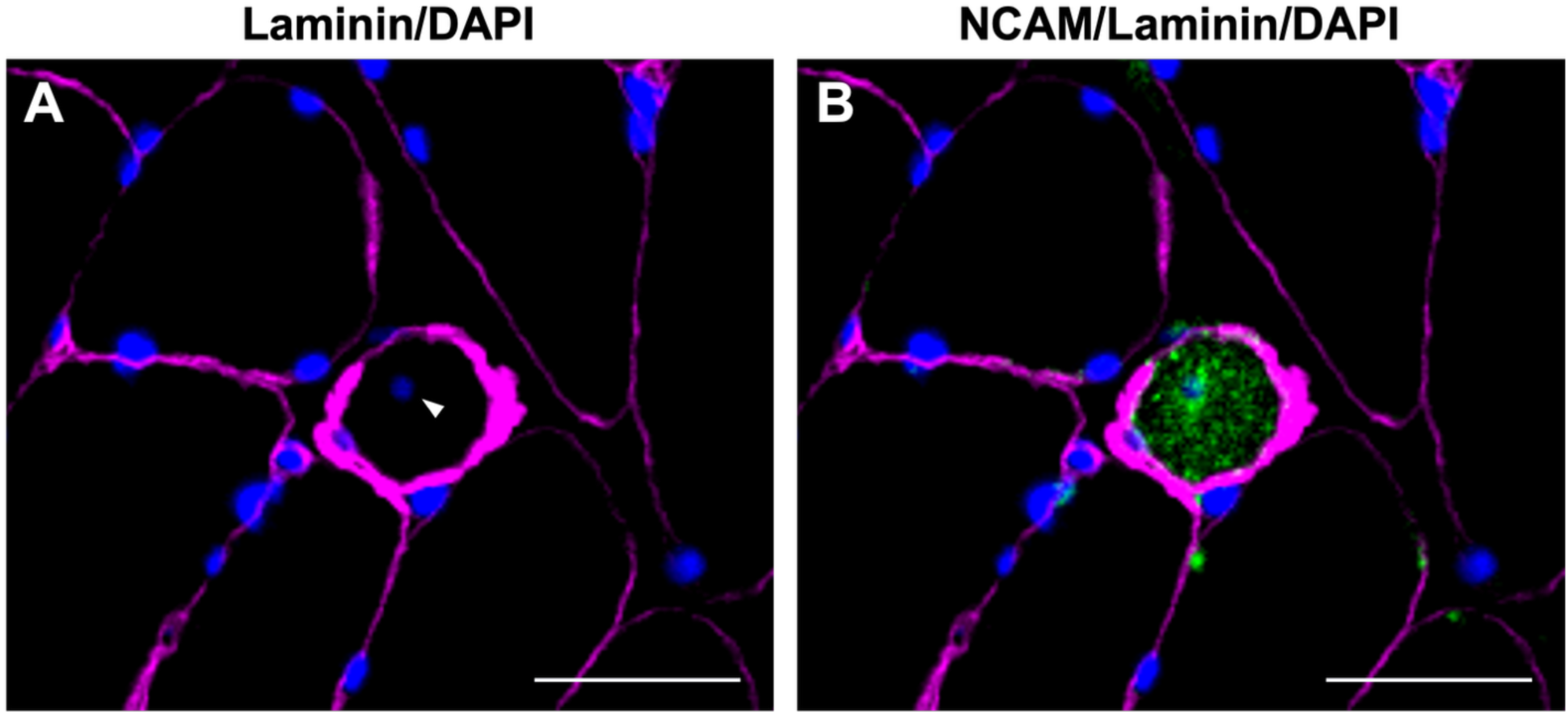
Representative NCAM+ Myofibre. *A)* Laminin (magenta) + DAPI (blue). DAPI staining shows the presence of a centrally located nucleus (white arrowhead). *B)* NCAM (green) + laminin (magenta) + DAPI (blue). One NCAM+ fibre is visible with a centrally located nucleus. Scale bar = 50 μm. NCAM, neural cell adhesion molecule.

## Discussion

Altered muscle morphology and cellular characteristics associated with knee OA may contribute to fibrillar muscle collagen accumulation and impaired muscle function. While the molecular response to acute resistance exercise appeared broadly preserved in knee OA, differences in MMP localization within the ECM highlight the importance of investigating exercise-induced changes at the compartmental level.

Acute resistance exercise increased gene expression of regulators of collagen breakdown (*MMP2, MMP9, MMP14, TIMP1,* and *TIMP2*) and enzyme activity potential of MMP2 and MMP9 in both volunteers with, and without moderate-to-severe knee OA. However, the increase in MMP area fraction, assessed by immunofluorescence, was blunted in muscle from volunteers with knee OA. Consistent with previous findings in muscle from volunteers with knee OA, greater muscle collagen was observed in volunteers with knee OA and occurred alongside reduced satellite cell abundance (Noehren et al., 2018; Serrão et al., 2014). The present study also demonstrated greater PDGFRα+ mesenchymal cell abundance in the muscle from volunteers with knee OA. PDGFRα+ mesenchymal cells positively correlated with fibrillar muscle collagen and negatively correlated with MMP9 area fraction at both the 0h and 72h time points, suggesting these cells may be involved in muscle collagen accumulation. Several measures of knee extensor strength were reduced in volunteers with knee OA and were negatively associated with fibrillar muscle collagen and SFI, providing further insight into potential mechanisms underlying muscle dysfunction in knee OA.

Consistent with other reports, the OA group demonstrated deficits in both isometric and isokinetic concentric knee extensor force compared with the control group (Noehren et al., 2018; Opara Zupančič & Šarabon, 2026). There is likely a combination of factors contributing to the strength deficits observed in knee OA. The knee OA group in the present study had smaller myofibre CSA, which may contribute to their reduced strength due to reduced contractile apparatus. However, no group differences in fibre type distribution (% fibres) were observed, and both groups had a greater proportion of their muscle CSA comprised of type I than type II myofibres. Thus, differences in fibre type distribution are unlikely to explain the lower strength observed in the OA group.

Another possible contributor to reduced muscle strength and quality in individuals with knee OA is neuromuscular function. The ECM plays a role in neuromuscular communication as the basal lamina at the neuromuscular junction, also referred to as the synaptic basal lamina, is a highly specialized structure with differential expression of laminin and collagen isoforms (Sanes et al., 1990). Consequently, alterations to the ECM surrounding the neuromuscular junction could influence neuromuscular transmission and muscle activation, though this was not measured in this present study and historically has been difficult to measure due to low success rates (∼17%) obtaining samples positive for NMJs (Aubertin-Leheudre et al., 2020).

Fibre denervation may contribute to strength loss during ageing (Soendenbroe et al., 2019) but did not present as a likely substantial contributor in the present study. Muscle denervation was assessed in the present study in a subset of participants (*n* = 10 control; *n* = 17 OA). NCAM is expressed on myofibres following denervation and is hypothesized to be involved in processes of reinnervation (Covault & Sanes, 1985). NCAM+ myofibres were identified in only four participants in the present study (*n* = 3 control; *n* = 1 OA). The low frequency of NCAM+ fibres in the present study suggests that there is minimal denervation in the present cohort of volunteers and is not likely a substantial contributor to reductions in muscle strength.

This contrasts another report in older adults in which NCAM+ fibres were observed in all participants in a cohort of healthy older males aged 65-94 years (*n* = 70), with 0.06-16.24% of fibres staining positive within individual participants (Soendenbroe et al., 2019). The participants from this study were originally recruited for a training study and thus were classified as sedentary or moderately active and performing no strength or other physical training daily (Heisterberg et al., 2018; Soendenbroe et al., 2019). In the subset of samples analyzed for NCAM+ staining in the present study, 80% of the controls and 76% of the volunteers with knee OA reported participation in weekly aerobic activity and weekly participation in lower-body resistance activity was 30% and 59% respectively. Mosole et al. (2014) demonstrated reduced myofibre denervation in muscles from seniors with a long history of high-level recreational sport participation compared to sedentary counterparts. Lifelong physical activity was not assessed in the present study, but the participants’ current physical activity levels may explain the low levels of denervation. Investigation of molecular indicators of myofibre denervation apart from NCAM (Soendenbroe et al., 2019) would further the understanding if myofibre denervation has a role in strength deficits in knee OA.

The greater collagen area observed in the OA group may contribute to impaired muscle function. Isokinetic peak torque was negatively correlated with percent collagen area in the present study and previous work has demonstrated isometric knee extensor strength to be negatively correlated with percent collagen area (Noehren et al., 2018). Losciale (2024) also demonstrated that eccentric isokinetic peak torque was nonlinearly related to ECM area in young adults. Increasing ECM content from 5-11% was related to increased torque, whereas past 13%, the relationship switched direction and more ECM content was related to reduced torque (Losciale, 2024). Increased collagen accumulation may impair force production by restricting myofibre radial expansion and lateral force transmission (Azizi et al., 2017; Spadoni et al., 2024).

Both the control and knee OA group demonstrated increased bulk gene expression (*MMP2, MMP9, MMP14, TIMP1* and *TIMP2)* and whole muscle MMP2/MMP9 enzyme activity potential 72h after acute resistance exercise. However, increased MMP2 and MMP9 protein area fractions were observed in only the control group. This apparent discrepancy between measures may reflect the complex regulation of MMPs at multiple levels including cell-specific transcription and translation, activation, inhibition, secretion and clearance (Cabral-Pacheco et al., 2020; Sternlicht & Werb, 2001).

MMP activity may be altered independently of gene expression or total protein content. For example, latent proenzymes can undergo activation, active MMPs can be inhibited by TIMPs, and changes in extracellular secretion can alter MMP actions in the ECM (Sternlicht & Werb, 2001). Like other contraction-induced changes in gene expression and protein abundance (Bishop et al., 2023), MMP gene expression, protein content and activity may be regulated independently of each other (Rullman et al., 2007). However, it is also possible that MMP regulation differs between the cell types responsible for their production and secretion. Myofibres, satellite cells, mesenchymal cells and immune cells express MMPs (Kedlian et al., 2024; Rayagiri et al., 2018; Rubenstein et al., 2025; Rullman et al., 2009). Gene expression and zymography analyses in the present study was from mixed muscle lysates; thus, compartment-specific changes may have been masked and complicates the interpretation of the mechanisms for altered MMP activity post-resistance exercise.

MMP2 and MMP9 area fractions were assessed by immunofluorescence, which provided information on MMP localization within the ECM. In contrast, MMP2 and MMP9 activity were assessed using gelatin substrate zymography which provided a mixed-muscle analysis. During zymography, proteins are denatured by SDS during electrophoresis and consequently, interactions between MMPs and endogenous inhibitors are disrupted (Snoek-van Beurden & Von den Hoff, 2005). Subsequent washing in Triton X-100 removes SDS and allows enzymes to be activated under the assay’s conditions. Latent proenzymes may autoactivate and contribute to the activity detected during the assay (Snoek-van Beurden & Von den Hoff, 2005). In the present study, the active and inactive MMP bands could not be discriminated on the zymogram, therefore, activity likely reflected the MMP activity potential within the tissue lysate rather than the activity that was physiologically present at the time of tissue collection (Snoek-van Beurden & Von den Hoff, 2005). Between-gel variability required zymography data to be expressed as fold changes from baseline, limiting comparisons of absolute MMP activity between groups. These methodological details provide context for interpreting the apparent discrepancy between MMP activity and MMP area fraction observed in the present study.

One explanation is that both groups similarly increased MMP protein following exercise but differed in its distribution. While MMPs are synthesized intracellularly (Sternlicht & Werb, 2001), muscle from volunteers with knee OA may exhibit delayed or reduced secretion in the ECM following exercise. MMPs located inside the myofibre may contribute to activity on the zymogram due to cell lysis during sample preparation but would not be detected by ECM-specific immunofluorescence. This would explain why increased MMP gene expression and activity was measured in OA muscle following exercise, but no changes in MMP area fraction were detected. This interpretation is consistent with the absence of baseline differences in MMP gene expression and area fraction between groups and suggests group differences arise during the post-translational processes governing MMP’s response to exercise.

Collectively, these findings highlight the importance of assessing MMPs at the gene expression, protein and enzyme activity levels. The findings demonstrated the importance of specifically investigating MMP protein localization, as the observed differences were undetected by whole-muscle analysis. Gene expression provides information about transcriptional responses, zymography provides an estimate of MMP activity potential, and immunofluorescence provides spatial information about MMP protein localization and abundance within the ECM. The divergence in findings between measures in the present study suggests that the impaired ECM response to acute resistance exercise in knee OA may occur downstream of transcription and may involve differences in MMP secretion or localization by differing cell types. However, it is possible that transcriptional differences between cell types may have gone unnoticed in the present study.

The present study identified several differences in myofibre morphology and cellular abundance between individuals with and without moderate-to-severe knee OA. The OA group had smaller myofibres and greater SFI, indicating greater myofibre shape irregularity. Previous reports have demonstrated that type I and type II myofibre SFI negatively correlated with peak isometric force (absolute and specific), as well as RTD (Soendenbroe et al., 2024). This is consistent with the present findings in which peak isometric peak force (absolute and specific) negatively correlated with myofibre SFI for both fibre types. Peak isokinetic force (absolute and specific) and RTD, however, only correlated significantly with type I fibres, though the sample size for the present analysis (*n* = 35) was much smaller than previous reports (*n* = 197) (Soendenbroe et al., 2024). Importantly, Soendenbroe et al. (2024) showed that myofibre shape could be improved following heavy resistance training. Although these findings were not obtained specifically in individuals with knee OA, they suggest that altered SFI may represent a modifiable feature of skeletal muscle and raise the possibility that resistance exercise could improve the structural characteristics of muscle in knee OA. However, mechanical constraint from the greater collagen content in OA muscle (Azizi et al., 2017; Spadoni et al., 2024) may reduce the adaptive potential of SFI to resistance training. Further research should investigate the mechanisms in which SFI is adaptable under different morphological conditions as well as how it may contribute to force generation.

The OA group had less satellite cells than the control group which is consistent with previous findings (Noehren et al., 2018). Neither group demonstrated satellite expansion in response to resistance exercise which contrasts a previous study in older males (Snijders et al., 2014). This is the only study, however, to our knowledge that has assessed satellite cell response in older adults 72h after acute resistance exercise (Dewi et al., 2023). Increases 24h and 48h after acute resistance exercise have consistently been demonstrated (Dreyer et al., 2006; McKay et al., 2012; Snijders et al., 2019; Walker et al., 2012), though only one of these reports assessed satellite cell changes in older females (Walker et al., 2012). The present study consisted of 61% females with an average age of 64 years. Thus, the lack of data in older females 72h after acute resistance exercise makes it possible that the absence of an increase observed in the present study reflects a true lack of satellite cell expansion and highlights an important area for future research. It is also possible, however, that the number of fibres assessed in the present study was not sensitive enough to distinguish exercise effects. Mackey et al. (2009) demonstrated that estimates of satellite cell content continued to improve with increasing numbers of fibres analysed. Previous satellite cell analysis in older males assessed approximately 100 type I and approximately 100 type II fibres for satellite cell analysis (Nederveen et al., 2015; Snijders et al., 2014), whereas the present study assessed approximately 65 per fibre type. Future studies should confirm the presence or absence of altered satellite cell response to exercise through assessment of a larger fibre number.

Compared with volunteers with knee OA, the control group demonstrated a greater fold change in satellite cell-MMP9 colocalization following acute resistance exercise. The blunted 72h increase in satellite cell-MMP9 colocalization in participants with knee OA may; therefore, indicate an altered satellite response to resistance exercise and impaired remodeling of the satellite cell niche. MMP9 colocalization with satellite cells following acute resistance exercise may reflect MMP9 production by satellite cells and subsequent secretion into the surrounding niche and niche remodeling by MMPs may be required for proper satellite cell activation and function. Rayagiri et al. (2018) demonstrated that activated but not inactivated satellite cells expressed MMP2 and MMP9 in cultured mouse single fibres. Rayagiri et al. (2018) also demonstrated a specific role for MMPs in basal lamina remodeling. Like MMPs, the laminin isoform, laminin-α1, was expressed from satellite cells only after activation and was localized to the satellite cell niche specifically. Inhibiting MMPs prevented laminin-α1 deposition in the satellite cell niche as well as transiently delayed satellite cell activation and permanently reduced satellite cell proliferation, self-renewal and differentiation (Rayagiri et al., 2018). Although these findings are from mouse muscle and satellite cells were autoactivated in cell culture rather than by exercise, they provide evidence linking MMPs with satellite cell niche and basal lamina remodeling as well as satellite cell function.

In the present study, when satellite cell counts from both groups were pooled, satellite cells were more likely to be MMP9+ after acute resistance exercise and incarcerated satellite cells were more likely to be MMP9+ than non-incarcerated cells. In a previous report, 12 weeks of combined resistance and interval training reduced satellite cell incarceration in older males (Nederveen et al., 2020). This suggests that satellite cell niche microstructure is adaptable to short-term exercise training and may be facilitated by MMP9. Further investigation of MMP expression and secretion specifically by satellite cells, immune cells and mesenchymal cells would help clarify the cellular contributions to ECM remodeling following resistance exercise. Analysis of satellite cell activation status after exercise in individuals with knee OA would also be valuable for understanding if differences in satellite cell function as well as abundance present.

The present study provides, to our knowledge, the first characterization of PDGFRα+ mesenchymal cells and capillaries in skeletal muscle from individuals with moderate-to-severe knee OA. No differences in capillaries per fibre were observed between groups, however, PDGFRα+ mesenchymal cells per mm² was greater in the OA group. Three cell types are generally recognized as contributors to skeletal muscle collagen accumulation: FAP cells, fibroblasts, and myofibroblasts (Joe et al., 2010). However, distinguishing among these populations can be challenging due to overlapping phenotypic characteristics. PDGFRα is commonly used to identify FAP cells in skeletal muscle (Joe et al., 2010; Uezumi et al., 2011), but this marker is not exclusive to this cell type. Connective tissue fibroblasts have also been shown to express PDGFRα, with evidence suggesting considerable overlap between FAP and fibroblast populations (Contreras et al., 2019; Murphy et al., 2011). Therefore, the PDGFRα+ mesenchymal cells identified in the present study may represent FAPs, fibroblasts, or a heterogeneous population of mesenchymal cells. The greater PDGFRα+ mesenchymal cell content in the OA group may contribute to the increased collagen accumulation observed in these individuals, as PDGFRα+ mesenchymal cells were positively correlated with percent collagen area. This hypothesis is supported by human data from ACL-injured and uninjured limbs as well as muscle from individuals with DMD (Cardone et al., 2023; Fry et al., 2017). Further characterization of the PDGFRα+ mesenchymal cell population in the present sample could provide insight into their role in muscle function and collagen accumulation. For example, Farup et al. (2021) identified a subpopulation of PDGFRα+ cells expressing CD90 that exhibited greater fibrotic potential, making this a marker of interest for future studies.

PDGFRα+ mesenchymal cells negatively correlated with MMP9 percent area both 0h and 72h following exercise, whereas no association was observed with MMP2. This may indicate a relationship between PDGFRα+ mesenchymal cells and MMP9-mediated ECM remodeling. However, the biological significance and directionality of this association remain unclear. Distinguishing between PDGFRα+ mesenchymal cell populations would provide further in insight into the nature of this relationship.

This study had several limitations that should be considered when interpreting the findings. First, the cross-sectional comparison between individuals with and without moderate-to-severe knee OA limits causal inference, such that observed associations cannot establish whether the identified alterations in skeletal muscle are a consequence of OA or contribute to its development. Furthermore, several of the mechanistic interpretations in the present study are based on associations between measures, which do not establish causality. Second, although multiple measures were used to assess MMPs, these measures do not directly quantify collagen degradation. MMPs represent one component of the complex processes regulating collagen turnover, and other proteolytic pathways, as well as collagen synthesis, may contribute to the greater collagen content observed in individuals with knee OA. Third, the post-exercise response was assessed at a single time point; therefore, earlier or later changes in MMP expression, activity, or cellular responses may have been missed. Our previous work demonstrated no changes in MMP or TIMP gene expression, or MMP protein abundance or activity, 6h after acute resistance exercise (Schweitzer et al., 2025), however, changes to MMP and TIMP gene expression as well as MMP activity have been demonstrated 24h after aerobic exercise (Rullman et al., 2009). Thus, assessments 24h and 48h post-exercise may provide additional insight into the temporal response to acute resistance exercise.

In conclusion, this study identified numerous alterations in muscle function, morphology, and cellular characteristics between volunteers with and without moderate-to-severe knee OA. Fibrillar collagen was negatively associated with knee extensor performance, supporting a potential role for altered ECM composition in strength deficits. Furthermore, the increase in PDGFRα+ mesenchymal cells and altered MMP protein response to acute resistance exercise provided potential cellular mechanisms contributing to collagen accumulation in knee OA. Importantly, changes to MMP transcription and activity potential in response to acute resistance exercise were preserved in OA muscle, whereas the increase in ECM-localized MMP protein (observed in controls) was blunted. This suggests that alterations in ECM remodeling may occur downstream of transcription in knee OA. Further characterization of PDGFRα+ mesenchymal cell populations and their fibrotic potential, alongside cell- and compartment-specific assessment of MMP and TIMP regulation, will be important for determining the mechanisms underlying collagen accumulation and altered muscle function in knee OA.

## References

Aubertin-Leheudre, M., Pion, C. H., Vallée, J., Marchand, S., Morais, J. A., Bélanger, M., & Robitaille, R. (2020). Improved human muscle biopsy method to study neuromuscular junction structure and functions with aging. The Journals of Gerontology: Series A, 75(11), 2098–2102. 10.1093/gerona/glz292

Azizi, E., Deslauriers, A. R., Holt, N. C., & Eaton, C. E. (2017). Resistance to radial expansion limits muscle strain and work. Biomechanics and modeling in mechanobiology, 16(5), 1633–1643. 10.1007/s10237-017-0909-3

Bellamy, N., Buchanan, W. W., Goldsmith, C. H., Campbell, J., & Stitt, L. W. (1988). Validation study of WOMAC: a health status instrument for measuring clinically important patient relevant outcomes to antirheumatic drug therapy in patients with osteoarthritis of the hip or knee. The Journal of rheumatology, 15(12), 1833–1840.

Binet, E. R., McKenna, C. F., Salvador, A. F., Martinez, I. G., Alamilla, R. A., Collao, N., Bodnariuc, G., Khan, N. A., Paluska, S. A., Burd, N. A., & De Lisio, M. (2023). Sex-based comparisons of muscle cellular adaptations after 10 weeks of progressive resistance training in middle-aged adults. Journal of Applied Physiology, 134(1), 116–129. 10.1152/japplphysiol.00274.2022

Bishop, D. J., Hoffman, N. J., Taylor, D. F., Saner, N. J., Lee, M. J., & Hawley, J. A. (2023). Discordant skeletal muscle gene and protein responses to exercise. Trends in Biochemical Sciences, 48(11), 927–936. 10.1016/j.tibs.2023.08.005

Callahan, D. M., Tourville, T. W., Miller, M. S., Hackett, S. B., Sharma, H., Cruickshank, N. C., Slauterbeck, J. R., Savage, P. D., Ades, P. A., Maughan, D. W., Beynnon, B. D., & Toth, M. J. (2015). Chronic disuse and skeletal muscle structure in older adults: sex-specific differences and relationships to contractile function. American Journal of Physiology-Cell Physiology, 308(11), C932–C943. 10.1152/ajpcell.00014.2015

Cabral-Pacheco, G. A., Garza-Veloz, I., Castruita-De la Rosa, C., Ramirez-Acuña, J. M., Perez-Romero, B. A., Guerrero-Rodriguez, J. F., Martinez-Avila, N., & Martinez-Fierro, M. L. (2020). The roles of matrix metalloproteinases and their inhibitors in human diseases. International journal of molecular sciences, 21(24), 9739. 10.3390/ijms21249739

Cardone, N., Taglietti, V., Baratto, S., Kefi, K., Periou, B., Gitiaux, C., Barneria, C., Lafuste, P., Pharm, F. L., Pharm, J. N., Panicucci, C., Desguerre, I., Bruno, C., Authier, F. J., Fiorillo, C., Relaix, F., & Malfatti, E. (2023). Myopathologic trajectory in Duchenne muscular dystrophy (DMD) reveals lack of regeneration due to senescence in satellite cells. Acta neuropathologica communications, 11, 167. 10.1186/s40478-023-01657-z

Contreras, O., Rossi, F. M., & Brandan, E. (2019). Adherent muscle connective tissue fibroblasts are phenotypically and biochemically equivalent to stromal fibro/adipogenic progenitors. Matrix Biology Plus, 2, 100006. 10.1016/j.mbplus.2019.04.003

Covault, J., & Sanes, J. R. (1985). Neural cell adhesion molecule (N-CAM) accumulates in denervated and paralyzed skeletal muscles. Proceedings of the National Academy of Sciences, 82(13), 4544–4548. 10.1073/pnas.82.13.4544

Dewi, L., Lin, Y. C., Nicholls, A., Condello, G., Huang, C. Y., & Kuo, C. H. (2023). Pax7+ Satellite Cells in Human Skeletal Muscle After Exercise: A Systematic Review and Meta-analysis: L. Dewi et al. Sports Medicine, 53(2), 457–480. 10.1007/s40279-022-01767-z

Dreyer, H. C., Blanco, C. E., Sattler, F. R., Schroeder, E. T., & Wiswell, R. A. (2006). Satellite cell numbers in young and older men 24 hours after eccentric exercise. Muscle & nerve: official journal of the American association of electrodiagnostic medicine, 33(2), 242–253. 10.1002/mus.20461

Farup, J., Just, J., de Paoli, F., Lin, L., Jensen, J. B., Billeskov, T., Roman, I. S., Cömert, C., Buch Moller, A., Madaro, L., Groppa, E., Göran Fred R., Kampmann, U., Gormsen, L. C., Pedersen, S. B., Bross, P., Stevnsner, T., Eldrup, N., Pers, T. N., & Jessen, N. (2021). Human skeletal muscle CD90+ fibro-adipogenic progenitors are associated with muscle degeneration in type 2 diabetic patients. Cell metabolism, 33(11), 2201–2214. 10.1016/j.cmet.2021.10.001

Fry, C. S., Johnson, D. L., Ireland, M. L., & Noehren, B. (2017). ACL injury reduces satellite cell abundance and promotes fibrogenic cell expansion within skeletal muscle. Journal of Orthopaedic Research, 35(9), 1876–1885. 10.1002/jor.23502

Hoffmann, M. D., Colley, R. C., Doyon, C. Y., Wong, S. L., Tomkinson, G. R., & Lang, J. J. (2019). Normative-referenced percentile values for physical fitness among Canadians. Health Rep, 30(10), 14–22. https://www.doi.org/10.25318/82-003-x201901000002-eng

Joe, A. W., Yi, L., Natarajan, A., Le Grand, F., So, L., Wang, J., Rudnicki, M. A., & Rossi, F. M. (2010). Muscle injury activates resident fibro/adipogenic progenitors that facilitate myogenesis. Nature cell biology, 12(2), 153–163. 10.1038/ncb2015

Kaminsky, L. A., Arena, R., Myers, J., Peterman, J. E., Bonikowske, A. R., Harber, M. P., Medina Inojosa, J. R., Lavie, C. J., & Squires, R. W. (2022). Updated reference standards for cardiorespiratory fitness measured with cardiopulmonary exercise testing: data from the Fitness Registry and the Importance of Exercise National Database (FRIEND). Mayo clinic proceedings, 97(2), 285–293. 10.1016/j.mayocp.2021.08.020

Kedlian, V. R., Wang, Y., Liu, T., Chen, X., Bolt, L., Tudor, C., Shen, Z., Fasouli, E. S., Prigmore, E., Kleshchevnikov, V., Pett, J. P., Li, T., Lawrence, J. E. G., Perera, S., Prete, M., Huang, N., Guo, Q., Zeng, X., Yang, L., & Zhang, H. (2024). Human skeletal muscle aging atlas. Nature aging, 4(5), 727–744. 10.1038/s43587-024-00613-3

Losciale, J. (2024). *Lost but not forgotten. An investigation of muscle performance after traumatic sport-related knee injuries* [Doctoral dissertation, University of British Columbia]. UBC Open Collections. 10.14288/1.0447249

Mackey, A. L., Kjaer, M., Charifi, N., Henriksson, J., Bojsen-Moller, J., Holm, L., & Kadi, F. (2009). Assessment of satellite cell number and activity status in human skeletal muscle biopsies. Muscle & Nerve: Official Journal of the American Association of Electrodiagnostic Medicine, 40(3), 455–465. 10.1002/mus.21369

McKay, B. R., Ogborn, D. I., Bellamy, L. M., Tarnopolsky, M. A., & Parise, G. (2012). Myostatin is associated with age-related human muscle stem cell dysfunction. The FASEB Journal, 26(6), 2509–2521. 10.1096/fj.11-198663

Mosole, S., Carraro, U., Kern, H., Loefler, S., Fruhmann, H., Vogelauer, M., Burggraf, S., Mayr, W., Krenn, M., Paternostro-Sluga, T., Hamar, D., Cvecka, J., Sedliak, M., Tirpakova, V., Sarabon, N., Musaró, A., Sandri, M., Protasi, F., Nori, A., & Zampieri, S. (2014). Long-term high-level exercise promotes muscle reinnervation with age. Journal of Neuropathology & Experimental Neurology, 73(4), 284–294. 10.1097/NEN.0000000000000032

Murphy, M. M., Lawson, J. A., Mathew, S. J., Hutcheson, D. A., & Kardon, G. (2011). Satellite cells, connective tissue fibroblasts and their interactions are crucial for muscle regeneration. Development, 138(17), 3625–3637. 10.1242/dev.064162

Nederveen, J. P., Joanisse, S., Séguin, C. M. L., Bell, K. E., Baker, S. K., Phillips, S. M., & Parise, G. (2015). The effect of exercise mode on the acute response of satellite cells in old men. Acta physiologica, 215(4), 177–190. 10.1111/apha.12601

Nederveen, J. P., Joanisse, S., Snijders, T., Ivankovic, V., Baker, S. K., Phillips, S. M., & Parise, G. (2016). Skeletal muscle satellite cells are located at a closer proximity to capillaries in healthy young compared with older men. Journal of cachexia, sarcopenia and muscle, 7(5), 547–554. 10.1002/jcsm.12105

Nederveen, J. P., Joanisse, S., Thomas, A. C., Snijders, T., Manta, K., Bell, K. E., Phillips, S. M., Kumbhare, D., & Parise, G. (2020). Age-related changes to the satellite cell niche are associated with reduced activation following exercise. The FASEB Journal, 34(7), 8975–8989. 10.1096/fj.201900787R

Noehren, B., Kosmac, K., Walton, R. G., Murach, K. A., Lyles, M. F., Loeser, R. F., Peterson, C.A., & Messier, S. P. (2018). Alterations in quadriceps muscle cellular and molecular properties in adults with moderate knee osteoarthritis. Osteoarthritis and Cartilage, 26(10), 1359–1368. 10.1016/j.joca.2018.05.011

Opara Zupančič, M., & Šarabon, N. (2026). Disease severity affects knee range of motion but not strength deficits in knee osteoarthritis: a systematic review and meta-analysis. Frontiers in medicine, 13, 1737973. 10.3389/fmed.2026.1737973

Rayagiri, S. S., Ranaldi, D., Raven, A., Mohamad Azhar, N. I. F., Lefebvre, O., Zammit, P. S., & Borycki, A. G. (2018). Basal lamina remodeling at the skeletal muscle stem cell niche mediates stem cell self-renewal. Nature communications, 9(1), 1075. 10.1038/s41467-018-03425-3

Roos, E. M., & Arden, N. K. (2016). Strategies for the prevention of knee osteoarthritis. Nature Reviews Rheumatology, 12(2), 92–101. 10.1038/nrrheum.2015.135

Rubenstein, A. B., Smith, G. R., Zhang, Z., Chen, X., Chambers, T. L., Ruf-Zamojski, F., Mendelev, N., Sze Cheng, W., Zamojski, M., Amper, M. A. S., Nair, V. D., Marderstein, A. R., Montgomery, S. B., Troyanskaya, O., G., Zaslavsky, E., Trappe, T., Trappe, S., & Sealfon, S. C. (2025). Integrated single-cell multiome analysis reveals muscle fiber-type gene regulatory circuitry modulated by endurance exercise. Genome Research, 35(7), 1664–1677. http://doi.10.1101/2023.09.26.558914

Rullman, E., Norrbom, J., Strömberg, A., Wågsäter, D., Rundqvist, H., Haas, T., & Gustafsson, T. (2009). Endurance exercise activates matrix metalloproteinases in human skeletal muscle. Journal of applied physiology, 106(3), 804–812. 10.1152/japplphysiol.90872.2008

Rullman, E., Rundqvist, H., Wagsater, D., Fischer, H., Eriksson, P., Sundberg, C. J., Jansson, E., & Gustafsson, T. (2007). A single bout of exercise activates matrix metalloproteinase in human skeletal muscle. Journal of applied physiology, 102(6), 2346–2351. 10.1152/japplphysiol.00822.2006

Sanes, J. R., Engvall, E., Butkowski, R., & Hunter, D. D. (1990). Molecular heterogeneity of basal laminae: isoforms of laminin and collagen IV at the neuromuscular junction and elsewhere. The Journal of cell biology, 111(4), 1685–1699. 10.1083/jcb.111.4.1685

Schweitzer, A. M., Koehle, M. S., Fliss, M. D., & Mitchell, C. J. (2025). Collagen remodeling increases after acute resistance exercise in healthy skeletal muscle irrespective of age. American Journal of Physiology-Cell Physiology. 10.1152/ajpcell.00992.2024

Serrão, P. R., Vasilceac, F. A., Gramani-Say, K., Lessi, G. C., BM Reiff, R., Mattiello-Sverzut, A. C., & Mattiello, S. M. (2014). Expression of receptors of advanced glycation end product (RAGE) and types I, III and IV collagen in the vastus lateralis muscle of men in early stages of knee osteoarthritis. Connective Tissue Research, 55(5-6), 331–338. 10.3109/03008207.2014.947368

Sharma, L. (2021). Osteoarthritis of the knee. New England Journal of Medicine, 384(1), 51–59. 10.1056/NEJMcp1903768

Snijders, T., Nederveen, J. P., Bell, K. E., Lau, S. W., Mazara, N., Kumbhare, D. A., Phillips, S. M., & Parise, G. (2019). Prolonged exercise training improves the acute type II muscle fibre satellite cell response in healthy older men. The Journal of physiology, 597(1), 105–119. 10.1113/JP276260

Snijders, T., Verdijk, L. B., Smeets, J. S., McKay, B. R., Senden, J. M., Hartgens, F., Parise, G., Greenhaff, P., & van Loon, L. J. (2014). The skeletal muscle satellite cell response to a single bout of resistance-type exercise is delayed with aging in men. Age, 36(4), 9699. 10.1007/s11357-014-9699-z

Snoek-van Beurden, P. A., & Von den Hoff, J. W. (2005). Zymographic techniques for the analysis of matrix metalloproteinases and their inhibitors. Biotechniques, 38(1), 73–83. 10.2144/05381RV01

Soendenbroe, C., Heisterberg, M. F., Schjerling, P., Karlsen, A., Kjaer, M., Andersen, J. L., & Mackey, A. L. (2019). Molecular indicators of denervation in aging human skeletal muscle. Muscle & Nerve, 60(4), 453–463. 10.1002/mus.26638

Soendenbroe, C., Karlsen, A., Svensson, R. B., Kjaer, M., Andersen, J. L., & Mackey, A. L. (2024). Marked irregular myofiber shape is a hallmark of human skeletal muscle ageing and is reversed by heavy resistance training. Journal of cachexia, sarcopenia and muscle, 15(1), 306–318. 10.1002/jcsm.13405

Spadoni, S., Todros, S., Reggiani, C., Marcucci, L., & Pavan, P. G. (2024). The role of the extracellular matrix in the reduction of lateral force transmission in muscle bundles: a finite element analysis. Computers in Biology and Medicine, 175, 108488. 10.1016/j.compbiomed.2024.108488

Stec, M. J., Graham, Z. A., Su, Q., Adler, C., Ni, M., Le Rouzic, V., Golann, D. R., Ferrera, P. J., Halasz, G., Sleeman, M. W., Lavin, K. M., Broderick, T. J., & Bamman, M. M. (2025). Combined endurance and resistance exercise training alters the spatial transcriptome of skeletal muscle in young adults. Iscience, 28(9). 10.1016/j.isci.2025.113301

Sternlicht, M. D., & Werb, Z. (2001). How matrix metalloproteinases regulate cell behavior. Annual review of cell and developmental biology, 17(1), 463–516. 10.1146/annurev.cellbio.17.1.463

Tarnopolsky, M. A., Pearce, E., Smith, K., & Lach, B. (2011). Suction-modified Bergström muscle biopsy technique: experience with 13,500 procedures. Muscle & nerve, 43(5), 716–725. 10.1002/mus.21945

Uezumi, A., Ito, T., Morikawa, D., Shimizu, N., Yoneda, T., Segawa, M., Yamaguchi, M., Ogawa, R., Matev, M. M., Miyagoe-Suzuki, Y., Takeda, S., Tsujikawa, K., Tsuchida, L., Yamamoto, H., & Fukada, S. I. (2011). Fibrosis and adipogenesis originate from a common mesenchymal progenitor in skeletal muscle. Journal of cell science, 124(21), 3654–3664. 10.1242/jcs.086629

Walker, D. K., Fry, C. S., Drummond, M. J., Dickinson, J. M., Timmerman, K. L., Gundermann, D. M., Jennings, K., Volpi, E., & Rasmussen, B. B. (2012). PAX7+ satellite cells in young and older adults following resistance exercise. Muscle & nerve, 46(1), 51–59. 10.1002/mus.23266

Zanotti, S., Gibertini, S., & Mora, M. (2010). Altered production of extra-cellular matrix components by muscle-derived Duchenne muscular dystrophy fibroblasts before and after TGF-β1 treatment. Cell and tissue research, 339(2), 397–410. 10.1007/s00441-009-0889-4

